# Structural Studies of an Anti-necroptosis Viral:Human Functional Hetero-amyloid M45:RIPK3 using SSNMR

**DOI:** 10.1101/2025.03.26.644761

**Authors:** Chengming He, Nikhil R Varghese, Eric G Keeler, Chi L L Pham, Brayden Williams, Stephan Tetter, Crystal Semaan, Karyn L Wilde, Simon H J Brown, James C Bouwer, Yann Gambin, Emma Sierecki, Megan Steain, Margaret Sunde, Ann E McDermott

## Abstract

The formation of RIP-homotypic interaction motif (RHIM)-based heteromeric amyloid assemblies between effector proteins such as RIPK1, ZBP1, or TRIF and the kinase RIPK3 serve as regulating signals for the necroptosis process, a key element of innate immune defense. Murine cytomegalovirus (MCMV) expresses the M45-encoded viral inhibitor of RIP activation (vIRA) which inhibits necroptosis in a RHIM-dependent manner. A pivotal question is how viral M45 forms hetero-amyloids with RIPK3 to effectively create an inhibitory assembly. We report a novel high-resolution structure of the M45:RIPK3 complex where M45 and RIPK3 alternately stack in an amyloid-state structure. Mutagenesis of the residues flanking the IQIG tetrad in M45 results in specific impacts on co-assembly with RIPK3, indicating an extended interface in the heteromeric fibrils. Other key interactions support the formation of stable viral:host fibrils. The M45: RIPK3 hetero-amyloid is likely to act as an anti-necroptotic signal by competing with formation of other pro-necroptotic species and introducing a barrier to RIPK3 autophosphorylation.

**Significance Statement:** This study investigates the structural biology of the necroptotic pathway, an understudied programmed cell death mechanism that plays a crucial role in innate immunity and has implications for infectious diseases, cell cycle regulation, and cancer. We present the high-resolution structure of a cross-species hetero-amyloid in which M45, a murine cytomegalovirus (MCMV) protein, co-assembles with human RIPK3 to inhibit necroptosis by competing with pro-necroptotic amyloids. Using solid-state NMR, cryo-EM, mutagenesis, and biophysical analyses, we uncover a novel structural paradigm for cross-species hetero-amyloids, shedding light on viral strategies to manipulate host immunity and protein interactions.

## Introduction

Necroptosis is a caspase-8-independent form of programmed cell death with essential roles in the promotion of tissue repair and the detection of pathogens. The best characterized trigger for necroptosis is activation of the tumor necrosis factor receptor (TNFR1) by TNF under conditions where caspase-8 is inhibited [1]. In this case, TNF does not initiate the formation of apoptotic complexes but instead a cytosolic complex comprising the receptor interacting protein kinases 1 and 3 (RIPK1 and RIPK3) is formed, known as the necrosome [2]. The interaction of RIPK1 and RIPK3 is facilitated by their RIP homotypic interaction motifs (RHIMs) [3-5]. RHIMs, approximately 18 amino acids in length, exhibit high sequence homology particularly in their conserved core tetrads [6], as shown in the sequence alignment in **Fig. 1(a)**. The necrosome, characterized as a functional hetero-amyloid, is activated through autophosphorylation and cross-phosphorylation [7]. Subsequent phosphorylation of mixed lineage kinase domain-like (MLKL) by activated RIPK3 results in conformational change and tetramerization of MLKL [8]. The oligomerized MLKL relocates to the plasma membrane, leading to cell membrane permeabilization and cell death [1, 9]. Necroptosis can be initiated by death receptors other than TNFR[1], TLR3 and TLR4 can induce necroptosis by forming a non-canonical necrosome through the RHIM-containing protein TIR-domain-containing adapter-inducing interferon β (TRIF) [10, 11] and another RHIM-containing protein, Z-DNA Binding Protein 1 (ZBP1, also known as DNA-induced activator of interferon, DAI), can initiate necroptosis in response to Z-form nucleic acid and viral infections [12].

**Figure 1.**
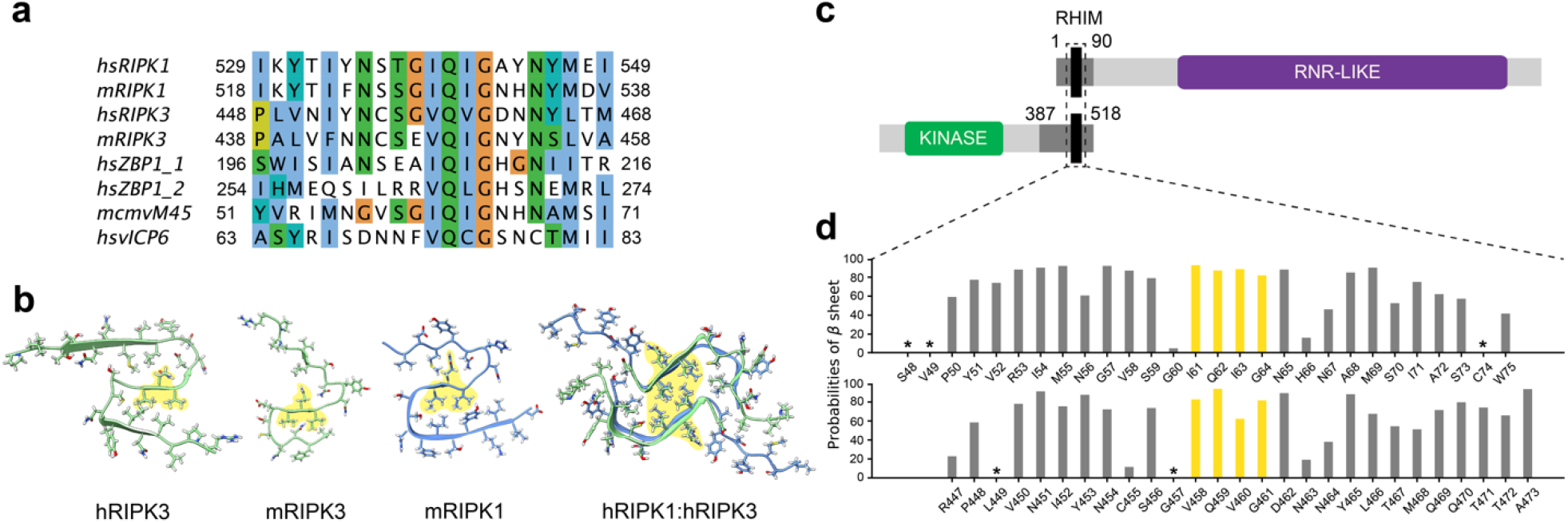
RHIM sequences, structural features, and protein context with β-Sheet probabilities of assigned residues. (a) Sequence alignment of RHIMs in human, mouse and viral proteins. (b) Single layer of RHIM core fibril structures by SSNMR with conserved tetrad shaded in yellow, from left to right: human RIPK3(7dac), mouse RIPK3(6jpd), mouse RIPK1(8ib0), and human RIPK1:RIPK3(5v7z). (c) Schematic of M45 and RIPK3 proteins, illustrating location of RHIMs relative to globular domains. The M45_1-90_ and RIPK3_387-518_ NMR constructs are colored in dark gray and the assigned regions in black. (d) β-sheet probabilities of M45 and RIPK3, inferred from Cα-Cβ or Cα-C’ chemical shifts. Asterisks indicate a failure of inference due to either a lack of sufficient data or unconventional chemical shifts. The core tetrad regions are highlighted in yellow, with two major interruptions in the β-sheets of M45 at positions G60 and H66, and in RIPK3 at positions G457 and N463.

RHIM-based interactions play crucial roles in development and adult tissue homeostasis [13, 14], protecting the host from pathogens and other challenges [15]. Moreover, aberrant regulation of key necroptosis players is associated with certain cancers [13] and dysregulation of this pro-inflammatory form of cell death has been associated with ischemic reperfusion injury and gut, lung and skin inflammatory pathologies [16, 17].

Elucidation of key RHIM-containing amyloid core structures is pivotal for comprehending the mechanisms underlying the induction and regulation of necroptosis, and RHIM-based interactions in inflammatory signaling. The structure of the necrosome core RIPK1:RIPK3 complex was determined using SSNMR [18], confirming the formation of a hetero-amyloid, wherein RIPK1 and RIPK3 are alternately packed and exhibit a two-fold symmetry around the fibril axis (**Fig. 1(b)**). The analysis unveiled a compact hydrophobic interface, incorporating the RHIM core tetrad, as well as Tyr stacking, and Gln, Asn, Thr/Ser, and Ser/Cys ladders, which collectively contribute to the stabilization of the structure. Subsequently, SSNMR structures of homomeric-RIPK3 and RIPK1 fibrils were reported [19-21] and reveal that the homomeric RHIM fibrils from mouse or human proteins share a serpentine amyloid core fold, with only one molecule per layer along the fibril long axis **(Fig. 1(b))**. A cryo-EM structure of human RIPK3 (PDB: 7da4) has also been reported, showing consistency with the SSNMR structure (PDB: 7dac) [20].

Within the context of pathogen infections, necroptosis assumes a critical role in the innate immune response [22]. To circumvent host defenses, pathogens have evolved several counteractive strategies aimed at inhibiting multiple cell death pathways [6]. For instance, Murine cytomegalovirus (MCMV), a betaherpesvirus, encodes suppressors of apoptosis as well as the M45-encoded viral inhibitor of RIP activation (vIRA) that blocks RHIM signaling and recruitment and phosphorylation of RIPK3 to prevent necroptosis [12, 23]. M45-mediated inhibition of necroptosis is essential for MCMV pathogenesis in mice [23]. The M45 protein features a RHIM at its N-terminus and includes an enzymatically inactive homolog of the large subunit (R1) of ribonucleotide reductase (RNR) [24] as depicted in **Fig. 1(c)**. Despite MCMV being a species-specific murine virus, the M45 protein can also disrupt necroptotic pathways in human cells by impeding the interactions of host RHIM-containing proteins. Our previous in vitro studies demonstrated the formation of hetero-amyloid fibrils containing M45 and RIPK3, facilitated by interaction between their respective RHIMs [25]. Indeed, M45:RIPK3 fibrils preferentially assemble over those containing RIPK1 and RIPK3, when all three proteins are mixed, suggesting a strategy by pathogens to mimic the hetero-amyloid motif used by their host and form an alternative structure that effectively serves as a blockade against necroptosis. This observation would also suggest a similarity in the interaction of other viral RHIMs, such as those in herpes simplex virus and varicella zoster, with host proteins [6, 24, 25]. The mechanism of viral inhibition of mammalian necroptosis could provide guidance for therapeutic disruption of unwanted or uncontrolled necroptosis [26].

In the present study, we determined the high-resolution structure of the viral:human M45:RIPK3 hetero-amyloid using SSNMR and validated it using biochemical and biophysical approaches. We discuss these findings in the context of how RHIM-mediated interactions, including viral protein containing complexes, regulate necroptotic pathways.

## Results

### M45 and RIPK3 Form 3 Segments of β-Sheet in RHIMs

The 90-residue N-terminal region of M45 (M45_1-90_) which includes the RHIM is sufficient to protect against TNF-induced necroptosis in human cells and spontaneously forms a hetero-amyloid with RIPK3_387-518_ in vitro [1]. **Fig.1(c)** and **Fig. S1** depict the protein constructs used in this study to prepare RHIM amyloid fibril samples. Partial uniform isotopic labeling schemes, illustrated in the top panel of **Fig. S2**, where only M45 or RIPK3 is uniformly labeled in the hetero-amyloid, were used to aid in the assignment of NMR peaks to residues of M45 and RIPK3 and to reduce the detection of intermolecular contacts. A total of 28 and 27 residues were unambiguously assigned in the NMR experiments for M45 and RIPK3, respectively, some of which are demonstrated in **Fig. S3**. Further details of the NMR resonance assignments can be found in the Methods section and the supplementary information. Residues N56 to N65 in M45 and C455 to D462 in RIPK3 were determined via the 3D backbone walk experiments with additional resonance assignments from 2D carbon homonuclear and carbon-nitrogen heteronuclear experiments (**Fig. S4**). The majority of the assigned regions were found to have a high β-sheet probability as determined from the assigned chemical shifts via PLUQin [27], as shown in **Fig.1(d)** and **Table S1**. Two major interruptions, occurring before and after the I(V)QI(V)G core tetrad in both proteins, indicates the presence of three β-sheet segments.

### Intramolecular Long-Range Contacts Reveal an ‘S’-shaped Core

We assigned 13 medium-range (1<|i-j|<4 residues) and 8 long-range (|i-j| > 4 residues) unambiguous intramolecular contacts in M45 and 11 medium-range and 10 long-range unambiguous intramolecular contacts in RIPK3 (**Fig.S5(a)** and **(b)** and **Table S2**). To determine these contacts, we employed partial sparse labeling schemes, illustrated on the bottom left of **Fig. S2**, where only M45 or RIPK3 is labeled with 1,3-^13^C-glycerol or 2-^13^C-glycerol, a common approach in structural characterization by SSNMR, to reduce the effect of dipolar truncation [28]. **Fig.2 (b)** and **(c)** show the reported contacts in a schematic backbone representation. The observed medium and long-range contacts were located exclusively within the RHIM domains of the proteins, supporting the existence of an ‘S’-shaped core, aligning well with the structural model previously reported by Liu and coworkers [20]. The long-range contacts were concentrated in the second turn of the ‘S’-shaped structure for both proteins. Further low-ambiguity contacts for both M45 and RIPK3 were also observed and can be found in **Table S3**.

### Intermolecular M45-RIPK3 Contacts Support the Formation of Hetero-Amyloid

To elucidate intermolecular contacts of the hetero-amyloid, we used additional mixed sparse labeling schemes, illustrated on the bottom right of **Fig. S2** (Details in Methods section). The NMR data showed a total of 9 unambiguous intermolecular contacts with 7 being found using long mixing carbon homonuclear correlation experiments and an additional 2 from carbon-nitrogen heteronuclear experiments. **Fig. 2(a)** and **S5(c)** depict the M45 intramolecular, RIPK3 intramolecular, and M45-RIPK3 intermolecular contacts in the DARR and TEDOR experiments, respectively. All intermolecular contacts are shown in **Fig. 2(d)** and summarized in **Table S2**. Evidence of the interaction between the RHIM core tetrads is bolstered by the four intermolecular contacts found here. The intermolecular contacts determined from these experiments strongly indicate the formation of a hetero-amyloid by M45 and RIPK3, which adopts parallel β-sheet registration.

**Figure 2.**
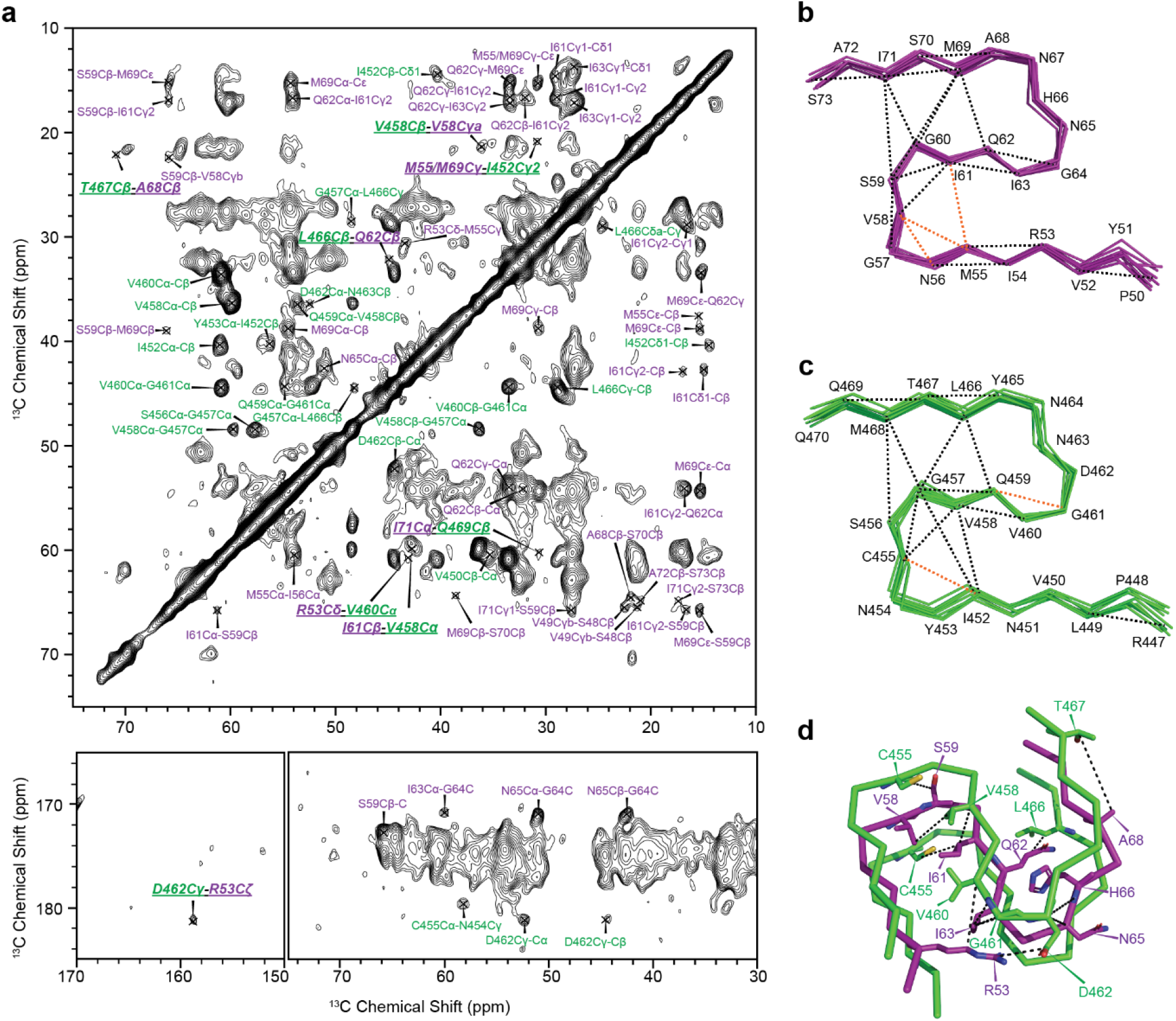
Spectra of 1,3-M45:2-RIPK3 and contacts summaries. (a) 500ms DARR spectrum of 1,3-^13^C-glycerol,^15^N-M45 : 2-^13^C-glycerol, ^15^N-M45. M45 and RIPK3 intramolecular contacts are labeled in purple and green, respectively. M45-RIPK3 intermolecular contacts are bold and underlined. (b), (c), and (d) represent the observed M45 intramolecular, RIPK3 intramolecular, and M45-RIPK3 intermolecular contacts mapped onto the structures, respectively. Unambiguous cross peaks are represented by black dashes, while ambiguous cross peaks are represented by orange dashes.

Three of the intermolecular contacts stand out as being separated by three or more residues in sequence alignment, I61Cβ(M45)-C455Cβ(RIPK3), V58Cγ2(M45)-V458Cβ(RIPK3), and R53Cζ(M45)-D462Cγ(RIPK3). These intermolecular contacts likely play crucial roles in the stability of the hetero-amyloid core structure.

### Calculation of M45-RIPK3 Hetero-Amyloid Structure

Structures were calculated based on the restraints listed in **Table S11** using simulated annealing with Xplor-NIH [29]. Motivated by the analogous structure of RIPK1:RIPK3 and the strong evidence from the NMR data of an ‘S’-shaped core, we modeled a representative unit of the quasi-infinite array hetero-amyloid with an alternating packing scheme of four copies each of the two proteins, as shown in scheme I of **Fig. 3**. To avoid biasing our conclusions, we also considered two additional schemes (II and III), one where M45 and RIPK3 choose their neighbors more randomly, instead of the strict alternating packing arrangement, and a second one where homo-M45 and homo-RIPK3 amyloids pack horizontally, as depicted in **Fig. 3**. Scheme II resulted in a configuration where segments of homo-M45 stack vertically with segments of homo-RIPK3. To ensure identical structural conformation in each instance of M45 and RIPK3, non-cyclization symmetry constraints were enforced separately on M45 and RIPK3 (excluding I452) using PosDiffPot terms in Xplor-NIH. In our calculations, we incorporated all three schemes, subjecting them to two rounds of annealing in torsion angle space (additional details of structural calculations in the Methods section).

**Figure 3.**
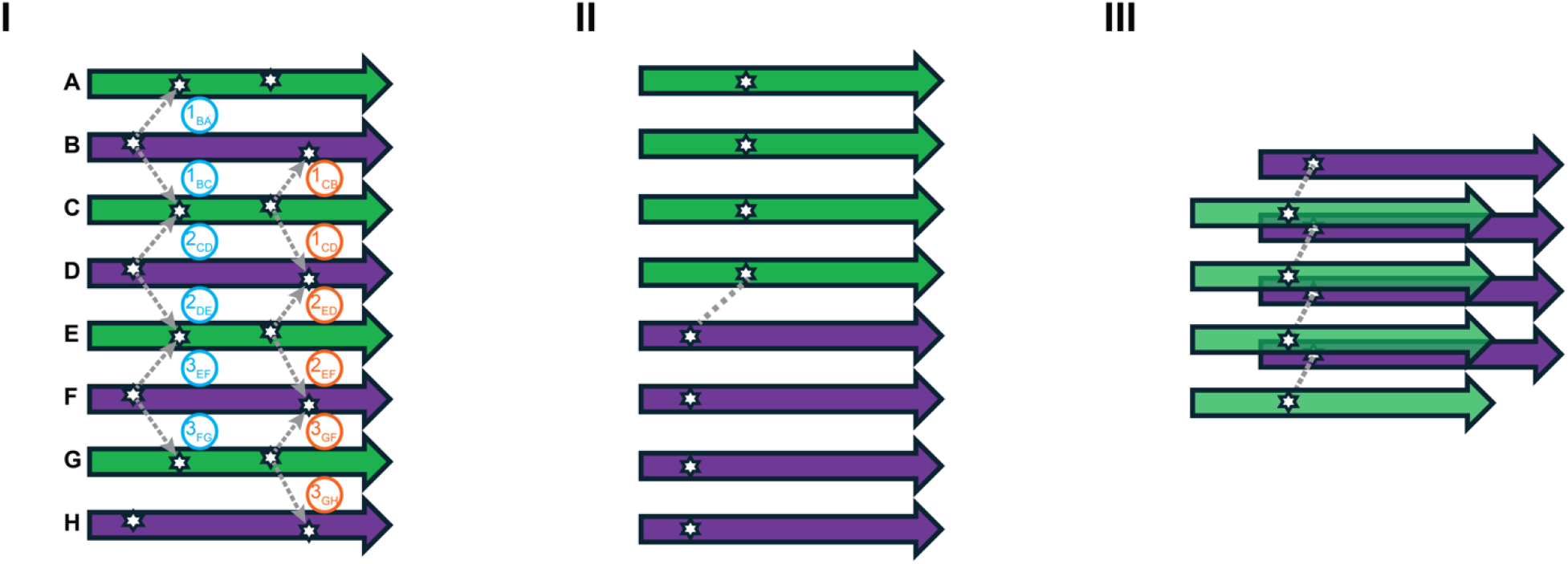
Arrangement of M45 and RIPK3 in Calculations. Scheme I represents the alternating stacking scheme between M45 (purple) and RIPK3 (green), with intermolecular contacts depicted as ambiguous distance constraints. Three sets of constraints are applied to each intermolecular contact, and scheme I specifies two constraints assignment, treating M45 as unambiguous (colored in blue) and treating RIPK3 as unambiguous (colored in orange). Scheme II represents the vertical packing of M45 and RIPK3 homo-amyloid. Scheme III represents the horizontal packing of M45 and RIPK3 homo-amyloid.

Within the context of scheme I, we established intermolecular contacts between M45 and RIPK3 as ambiguous distance constraints. This ambiguity stemmed from the structural arrangement that allowed a specific atom in M45 to interact with a second specific atom from RIPK3 both in the hetero interface surface above or in another RIPK3 copy in the distinct surface below, denoted as BA and BC contacts (illustrated by the 1BA and 1BC dotted lines in **Fig. 3** scheme I), respectively. We term such ambiguity “symmetric ladder ambiguity” to distinguish it from ambiguity due to chemical shift overall. Within scheme I (tetramer of heterodimers), three or four equivalent examples of each of the possible interaction will appear; in other words the tetramer is large enough to represent all interactions. Since generally only one of these two interactions is close enough to be NMR detected, we treated the interaction as either occurring with the hetero partner above or with the one below or both, and we employed low-ambiguity distance constraints. Scheme I of **Fig. 3** presents two distinct approaches for assigning constraints with “symmetric ladder ambiguity”. In one approach, the M45 atom was considered unambiguous, while that in RIPK3 was treated as ambiguous (indicated in blue). Consequently, atoms in the copy of M45 at one edge (segment H) remained unconstrained by intermolecular contacts, while remaining constrained by the imposed symmetry. The alternative approach involved considering RIPK3 as unambiguous and M45 as ambiguous (indicated in orange), with analogous end effects. In scheme II, each intermolecular contact was uniquely assigned, as there was only one interface. In scheme III, intermolecular contacts were presumed to occur at every layer.

In Scheme I, we observed two ensembles of structures in the first round of annealing, with a major species comprising around 75% and a minor species around 25% (as shown in **Fig. S9**). This divergence originates from ambiguities in the inter-layer arrangement of M45:RIPK3 intermolecular contacts, particularly the R53 (M45)-D462 (RIPK3) interaction. Incorporating low-ambiguity cross-peaks in the second round of annealing caused the output structures to converge to the major species.

Following similar computation procedures on schemes II and III, we found that scheme II converged to an ‘S’-shaped core as well. The intensity of the intermolecular M45:RIPK3 cross peaks in the experimental NMR data disfavors scheme II as it would predict relatively sparse and weak intermolecular contacts. Consequently, the structures from scheme II were not included in further analyses. When using scheme III, the calculations did not converge under the specified conditions. For the lowest-energy structure, shown in **Fig. S11(a)**, substantial violations remain in all constraints, particularly in inter- and intramolecular distances and dihedral angles. Meanwhile, the RMSD distribution of the lowest 160 structures relative to the best structure shows a broad distribution with large RMSDs (**Fig. S11(b)**), in contrast to the narrow distribution in **Fig. S10(c)** and **Fig. S12(c)** for Scheme I.

### Structural features of M45:RIPK3 hetero-amyloid core

We report 10 structures of M45:RIPK3 with RMSD among core backbone heavy atom being 0.52Å; core residues include 52-73 for M45 and 449-470 for RIPK3. The resulting structural model of the M45:RIPK3 hetero-amyloid, as shown in **Fig. 4**, exhibits an ‘S’-shape on both M45 and RIPK3, each featuring a hydrophobic core around the initial turn comprising eight residues: M55, V58, I61, and I63 in M45; and I452, C455, V458, and V460 in RIPK3. This hydrophobic core contains two crucial experimental intermolecular contact restraints, I61Cβ(M45)-C455Cβ(RIPK3) and V58Cγ2(M45)-V458Cβ(RIPK3). The structural integrity of this double turn structure is maintained through several features: two key sets hydrogen bonds, colocalized polar residue ladders, and colocalized hydroxy group-containing side chains. The interaction between R53 on M45 and D462 on RIPK3, experimentally evident in the intermolecular contact between R53Cζ(M45) and D462Cγ(RIPK3), forms an anchoring salt-bridge which tethers the N-terminal tail of M45 to the second strand of RIPK3. Another hydrogen bond set involves the side chains of Q62 in M45 and Q459 in RIPK3, along with the backbones of A68 in M45 and Y465 in RIPK3, supporting the second turn. Colocalized polar residues, like Q62 and Q459, form ladders along the fibril axis, through hydrogen bonding networks, reinforcing the cross-β structure. An additional hydrogen bond network formed by H66 in M45 and N463 in RIPK3, with their corresponding hydrogen bond acceptors on the backbone of the strands, enhances the stability of the second turn. A Ser-Thr ladder involving S70 in M45 and T467 in RIPK3 and a Ser-Ser ladder involving S59 in M45 and S456 in RIPK3 further support the stability of the structure via their colocalized hydroxy group-containing side chains.

**Figure 4.**
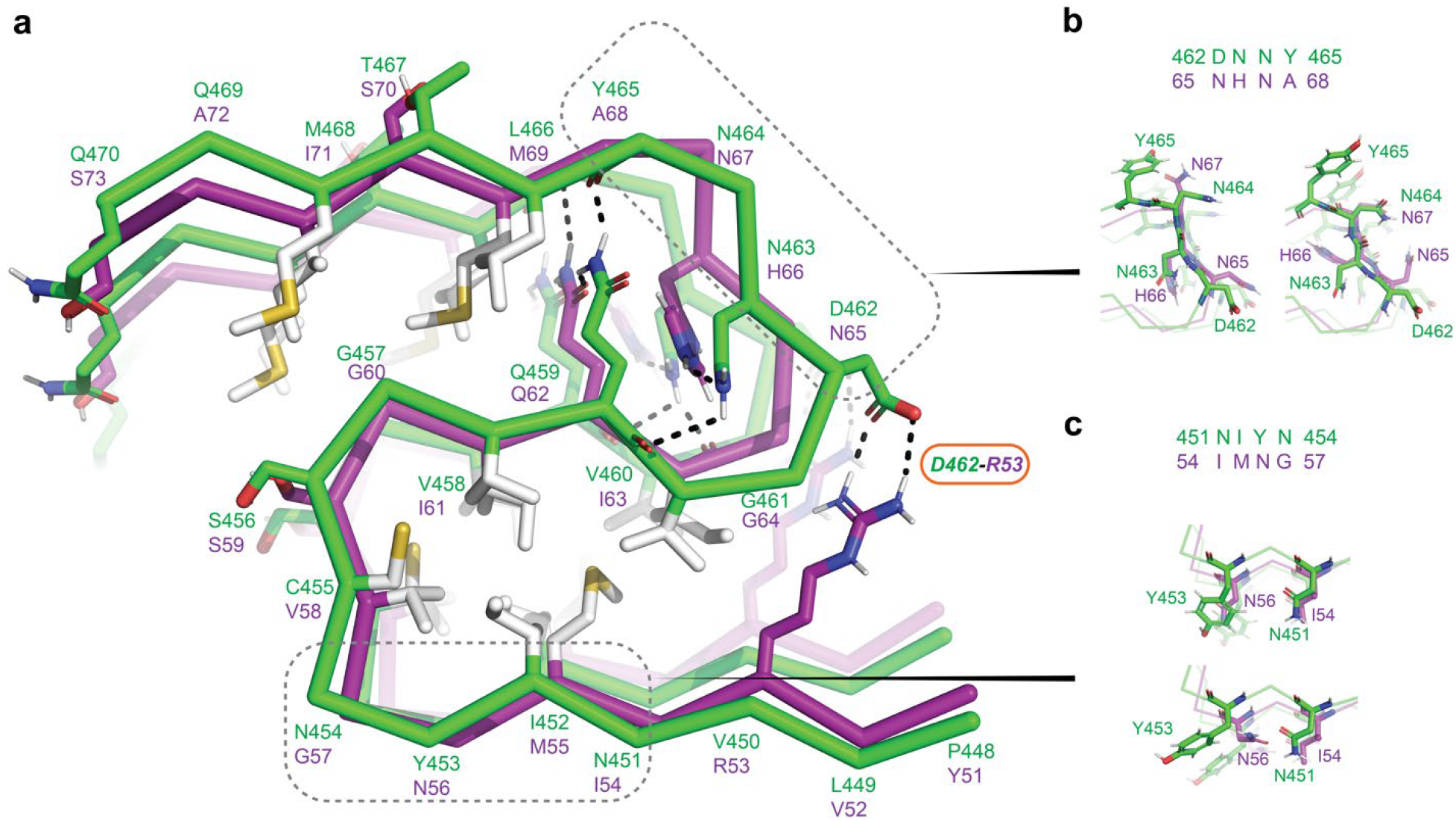
Structure of hetero-amyloid M45:RIPK3. (a) Ribbon diagrams of M45 (purple) and RIPK3 (green), with core hydrophobic residues and stabilizing polar residues highlighted. Hydrogen bonds are represented by black dashes, with the D462-R53 contact emphasized based on experimental data. Two sequence mismatches, upstream and downstream of the core tetrads, are highlighted. (b) and (c) show the sidechain variabilities of two brief sequence mismatches: D462-Y465/N65-A68 and N451-N454/I54-G57, respectively.

Despite the RHIM sequence similarities in M45 and RIPK3, residue mismatches lead to differences and heterogeneities among the calculated structural ensemble. **Fig. 4 (b)** and **(c)** illustrate two brief segments with sequence disparities resulting in two distinct configurations in the final structures. The first such segment comprises residues 54-57 (IMNG) on M45 and 451-454 (NIYN) on RIPK3. On the top of **Fig. 4(c)**, the first configuration features π-stacking between Y453 and N56 and formation of a hydrogen bond between the hydroxyl group of Y453 and the amine group of N56, while I54 and N451 are also stacked. The alternative arrangement involves hydrogen bonds between the side chains of N56 and N451, as depicted on the bottom of **Fig. 4(c)**. In the second example involving residues 65-68 (NHNA) of M45 and 462-465 (DNNY) of RIPK3, as illustrated in **Fig. 4(b)**, residues N65, H66, and N67 display variability in their side chains, interacting with adjacent or colocalized residues in RIPK3. A majority of the residues demonstrating this variability are oriented toward the solvent, suggesting that the M45:RIPK3 complex possesses a more rigid and well-defined core structure, while the solvent-exposed residues exhibit greater flexibility.

### Comparison between homo- and hetero-amyloids

We performed comparisons between the SSNMR data and the structural models of the novel RIPK3(M45) and RIPK3(homo) data and previous reported RIPK3 data in homomeric and heteromeric complexes. Here the notation RIPK3(M45) refers to RIPK3 in the M45:RIPK3 hetero-amyloid and RIPK3(homo) refers to RIPK3 in homomeric fibrils. With the exception of Q459Cδ in the core tetrad of RIPK3, our RIPK3(M45) SSNMR data demonstrate close agreement with previously reported RIPK3(homo) [20], however larger differences were found with RIPK3(RIPK1) [18]. **Fig. 5(a), S14(a)** and **(b)** show the comparison between the RIPK3(homo) and RIPK3(M45) spectra reported here. The carbon homonuclear correlation experiments indicate a high level of consistency, including two sets of isoleucine resonances in both samples, despite I452 being the only isoleucine in the RIPK3 construct. The cause of this peak doubling remains unclear. However, the intensity of the sets of I452 cross-peaks differs in the samples. The peak intensity of N463 is stronger in RIPK3(homo), which we attribute to the disruption of a more stable asparagine ladder in the RIPK3(homo) by H66 of M45 in the M45:RIPK3 complex. The Cγ-Cα peak of D462 reveals an upfield shift in RIPK3(M45). This agrees with a chemical shift change for a protonated carboxylic group [30] as indicated by the possible salt-bridge between R53Cζ (M45) and D462(RIPK3) (**Fig. 2(a)**). Notably, in the NCA spectra (**Fig. S14(b)**), N463 and A473 exhibit stronger signals in RIPK3(homo), whereas N451 shows some disorder in the RIPK3(M45) spectrum.

**Figure 5.**
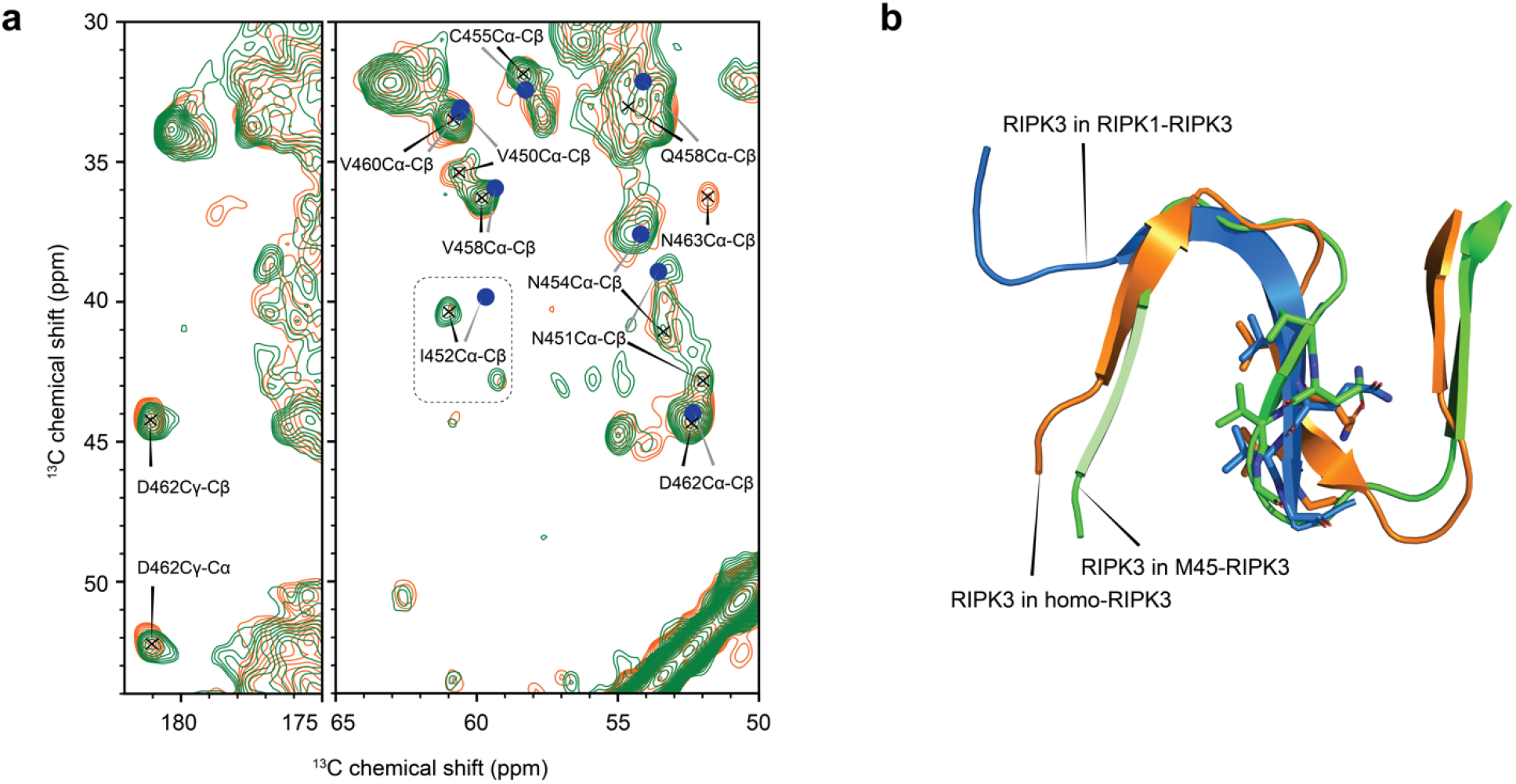
Spectral and structural comparisons of RIPK3 in three fibrillar complexes. (a) 50ms DARR spectra of RIPK3(homo) in orange and RIPK3(M45) in green. The Cα-Cβ cross peaks around the core tetrad of RIPK3(RIPK1) are represented by solid blue circles. (b) Structural alignment of RIPK3(homo) and RIPK3(RIPK3) to RIPK3(M45). Backbone heavy atoms of 450-469 in RIPK3(homo) and 450-462 in RIPK3(RIPK3) are aligned.

We aligned our structural model of RIPK3 with the previously reported RIPK3(homo) structures [20] and RIPK3(RIPK1) structure [18], as depicted in **Fig. 5(b)**. While the RIPK3(homo) SSNMR structural model exhibits considerable consistency to the structure of RIPK3 in the M45 complex, with a heavy atom backbone RMSD of 2.2 Å when aligning the core region (450-469) and a reduction to 1.5 Å when excluding the first β-strand and aligning residue 457-469 (as shown in **Fig. S14(c)**), there are several factors contributing to the differences between these structures. The M45:RIPK3 hetero-amyloid has a more compact hydrophobic core resulting from the shorter distance between the first and second strands in our model, attributed to the formation of the R53(M45)-D462(RIPK3) stabilizing salt-bridge. Moreover, while in our M45:RIPK3 model the β-strands are nearly coplanar, the first β-strand in the RIPK3(homo) model protrudes from the plane of the other two strands. With an RMSD of 1.8 Å and 1.3 Å for the two regional alignments, we found improved consistency with the cryo-EM structure (**Fig. S14 (c)**). On the other hand, RIPK3(RIPK1) possesses a shorter C-terminal tail compared to RIPK3(M45) and has a backbone RMSD of 2.3 Å when aligning the core (residues 450-462).

The carbon homonuclear correlation experiments are notably consistent with few differences observed when comparing M45(homo) and M45(RIPK3) (**Fig. S15(a)** and **(b)**). Two residues, I54 and M55, both near the N-terminus of the core, exhibit differences. The latter has stronger cross-peak intensities and improved resolution in M45(RIPK3) and there is a chemical shift change for I54Cγ2,Cβ. S59 and H66, both located at turns exhibited chemical shift changes in the NCA spectra (**Fig. S15(b)**). Additionally, several unassigned glycine peaks were only observed in the M45(RIPK3) sample. While the assigned glycine residues (57, 60, 64), located within the amyloid core, demonstrated similarities in the two samples, there is a cluster of glycine residues preceding the core (**Fig. S15(c)**). We propose the unassigned glycine resonances arise from these glycine residues, suggesting that the M45(RIPK3) amyloid possesses a longer structured N-terminal tail.

We also investigated homo-M45 amyloid fibrils by cryo-EM, selected for study because the hetero-amyloid presents technical challenges. M45 readily assembles into dense networks of homo-M45 fibrils in the absence of denaturants. Given that helical reconstruction requires isolated filaments, these networks pose an inherent hinderance to the helical reconstruction of homo-M45. Despite this limitation, a total of 1,194 filament segments were manually picked from 14,767 micrographs and facilitated the helical reconstruction of the homo-M45 amyloid core to a resolution of 3.96 Å (FSC 0.143, **Fig. 6(a)**). This reconstructed cryo-EM map features an S-shaped fold with an imposed left-handed twist of −5.34° and a helical rise of 4.8 Å (**Fig. 6(a)**). Modelling of the secondary structure of M45 subunits within the homo-M45 fibril core demonstrates the presence of at least two β-strands within the larger S-fold architecture, similar to that observed in the NMR structure of M45:RIPK3 (**Fig. 6(a)** inset). Notably, the cryo-EM map indicates that the central β-strand is more extended in homo-M45 fibrils, consistent with the chemical shift changes observed for S59 and H66. Overall, the fact that M45 can also form serpentine amyloids is consistent with the observation that it readily forms hetero-amyloids with RIPK3.

**Figure 6.**
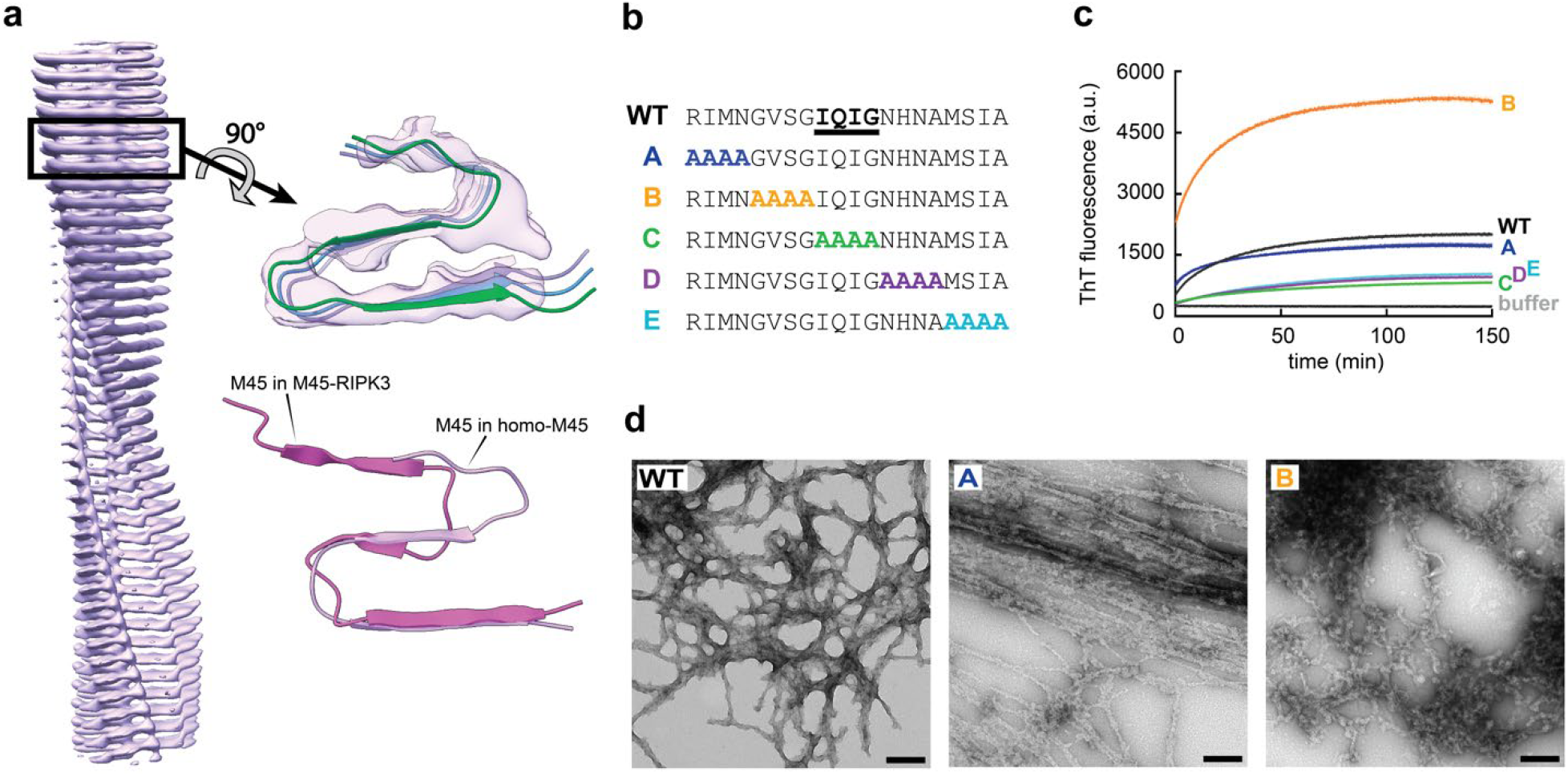
M45 homomeric amyloid assembly. (a) Cryo-electron density map from homomeric M45 fibrils. Insets show fibrillar cross-section with modelled secondary structure of M45 in homomeric fibrils and a comparison with the secondary structure of M45 in heteromeric M45:RIPK3 fibrils. (b) Sequences of WT M45 and AAAA mutants mutA–mutE. (c) Thioflavin T fluorescence assay of amyloid assembly by WT and AAA mutants. Fluorescence emission (arbitrary units) at 485 nm; average of triplicate values, error bars represent the range. (d) Negative stain transmission electron micrographs of assemblies formed by WT and the AAAA mutants of M45 for which fibrils were observed. Scale bar 200 nm.

### Biochemical and biophysical validation of M45:RIPK3 structure

The roles of specific interactions in the flanking sequence elements (discussed above in connection with our structure) were probed using alanine scanning methods. By introducing five AAAA substitutions to the M45 RHIM, spanning R53 to A72 (mutants A-E; **Fig. 6(b)**), mutagenesis confirmed the extent of the stabilizing homo-M45 core and the role of key stabilizing interactions in the M45:RIPK3 hetero-amyloid. First, the ability of these mutants to form homo-amyloid was investigated by Thioflavin T (ThT) fluorescence assays and negative stain transmission electron microscopy (TEM). WT M45 formed amyloid rapidly at pH 7.4 and the homomeric fibrils formed by the WT RHIM displayed inter-fibrillar contacts and form a network (**Fig. 6(c)** and **(d)**). AAAA substitutions in the N terminal region of the M45 RHIM had a noticeable impact on amyloid assembly. Mutant A displayed similar levels of ThT fluorescence to WT but formed fibrils that associated laterally. Substitution of the GVSG sequence by AAAA in mutant B resulted in a large increase in the intensity of ThT fluorescence, likely reflecting increased amyloidogenicity or local structural changes arising from this sequence change where two glycine residues which have low amyloid-forming potential are replaced by alanine residues. Mutant B formed numerous short and curvy fibrils. Changes to the core tetrad and to residues C-terminal to the IQIG sequence disrupted the ability of M45 to self-assemble into fibrils. Although a small increase in ThT fluorescence signal was detected for M45 mutants C, D and E, no fibrils were observed by TEM.

Single molecule fluorescence validated the role of key residues and intermolecular interactions identified by SSNMR and cryo-EM. Samples of mCherry-M45 and YPet-RIPK3, alone or as mixtures, were prepared at sub-micromolar concentration under assembly-permissive conditions and the fluorescence intensity was collected from a small detection volume, through which the proteins were freely diffusing. The amplitude and frequency of the fluctuations reflects the extent of assembly into multimeric species with a range of sizes. Thioflavin T fluorescence and TEM indicate these species are amyloid fibrils (Fig. 6 and S1d). WT M45, M45 mutant B and WT RIPK3 rapidly formed homo-amyloid when prepared individually under these conditions; M45 mutant A showed a very low level of self-assembly but no self-assembly of M45 mutants C, D or E was detected on this timescale (**Fig. 7(a)**). The ability to undergo heteromeric assembly was determined by Two-Colour Coincidence Detection from mixtures containing RIPK3, and WT or mutant M45 (**Fig. 7(b)** and **Fig. S16**). The formation of WT M45:RIPK3 hetero-amyloid fibrils resulted in strong and coincident mCherry and YPet fluorescence intensity from the two proteins, with 80% of oligomers formed containing both proteins. Loss of key intermolecular interactions due to alanine substitution of R53 (M45 mutant A), S59 (M45 mutant B), Q62 (M45 mutant C), H66 (M45 mutant D) and S70 (M45 mutant E) notably diminished the tendency for M45 to form hetero-amyloid with RIPK3. Although many multimers of M45 mutant B were detected in the mixture with RIPK3, the extent of coincidence of the YPet and mCherry fluorescence signals was reduced, indicating that the fibrils formed were predominantly homo-M45 and homo-RIPK3 amyloid.

**Figure 7.**
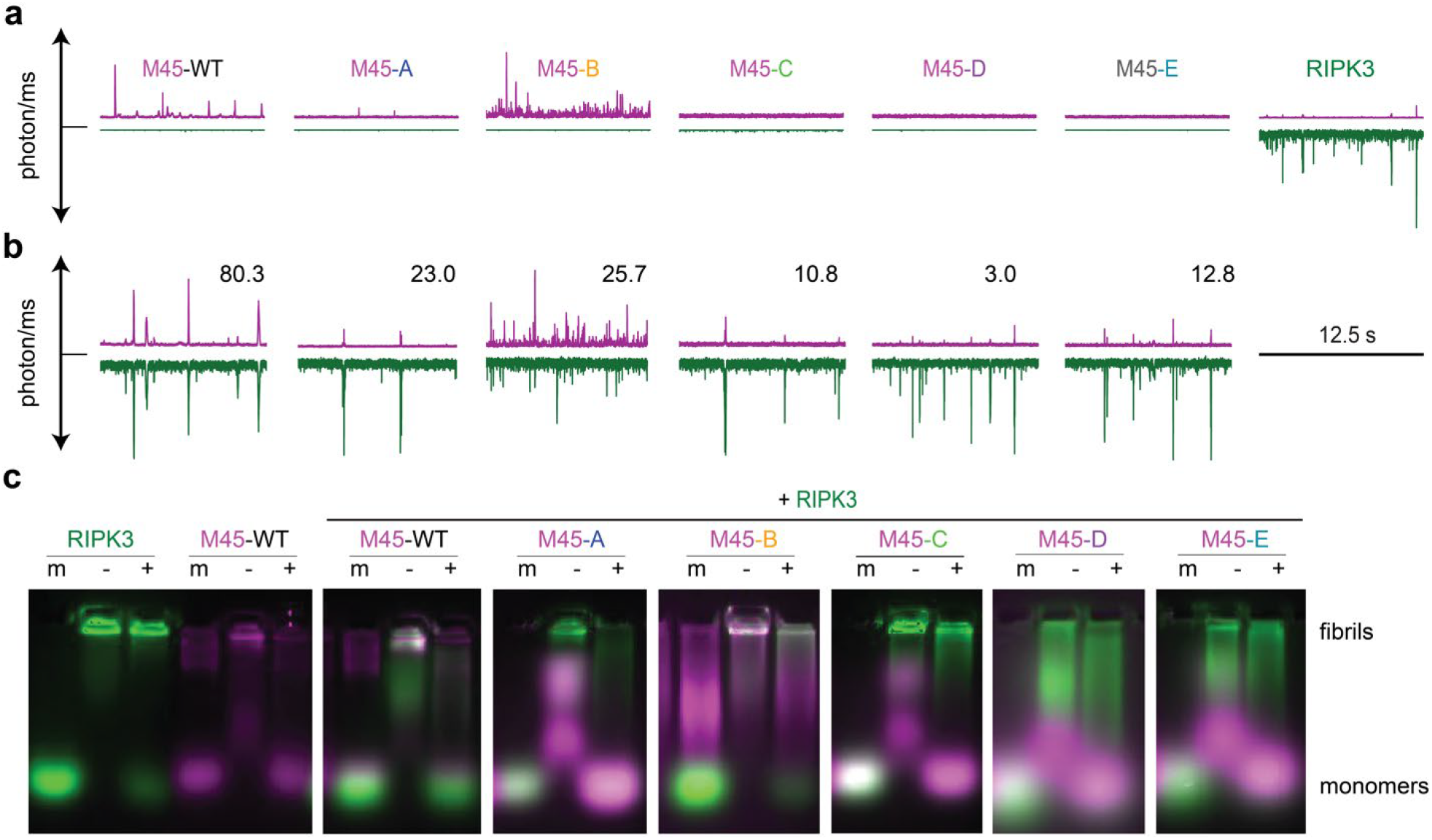
M45 homomeric assembly, and heteromeric assembly with RIPK3. (a) Simultaneous detection of mCherry (positive y-axis) and YPet (negative y-axis) fluorescence from homomeric samples of WT or AAAA mutants of mCherry-M45 or YPet-RIPK3. Samples contained only one type of protein but a small amount of bleed through into the mCherry channel occurred when very large YPet-RIPK3 assemblies passed through the confocal volume. (b) Simultaneous detection of fluorescence from mCherry-tagged WT and mutant M45 and YPet-RIPK3 in samples containing both RHIM proteins. The numbers in each panel represent the percentage of events where WT or mutant M45 and RIPK3 are detected in the same diffusing complex, over 180s, from full traces presented in **Fig. S11**. (c) SDS semi-denaturing agarose gel electrophoresis indicates the nature of species observed when RIPK3, and WT or AAAA mutants of M45, are maintained individually in urea (m), or incubated alone or together under assembly-permissive conditions (−) and then treated with 2% SDS (+).

The contribution of intra and intermolecular interactions to RHIM amyloid stability was probed by 2% SDS treatment of fibrils, analysed by semi-denaturing electrophoresis (**Fig. 7(c)**). RIPK3 homo-amyloid is relatively stable to 2% SDS treatment, while M45 homo-amyloid is partially dissociated to monomer. Strikingly, WT M45:RIPK3 hetero-amyloid is predominantly dissociated to monomer by 2% SDS, reflecting the interdigitation of the viral RHIM that disrupts the perfectly matched sidechain packing in RIPK3 homo-amyloid. Analysis of the assemblies formed by RIPK3 and the M45 mutants reflected the loss of stabilizing hetero-interactions. All five M45 mutants interacted with RIPK3 to some extent, as reflected in a change in the distribution of RIPK3, from all stable RIPK3 fibrils in the wells, to a distribution of smaller RIPK3 species migrating in the gel following incubation with 2% SDS. None of the M45 mutants was able to form SDS-resistant hetero-amyloids with RIPK3. The increased amyloidogenicity of M45 mutant B was reflected in the formation of homo-oligomers when diluted out of urea during electrophoresis. Although mCherry and YPet-positive fibrils formed in mixtures of mutant B and RIPK3, as seen in the two-colour coincidence experiments, these are predominantly homo-fibrils of either M45 or RIPK3, which are both relatively resistant to depolymerisation by 2% SDS.

M45 is a potent inhibitor of RHIM signalling in both human and murine cells, even though the mouse is the natural host for MCMV. The RHIM-based interactions between murine RIPK3 (mRIPK3) and M45 are similar to those between human RIPK3 (hRIPK3) and M45 (**Fig. S17**). The mRIPK3 RHIM forms homo-amyloid fibrils in vitro at a similar rate to the human RHIM and likewise shows rapid incorporation into hetero-amyloid structures with M45 (**Fig. S17(a)** and **(b)**). The incorporation of the viral M45 RHIM has a similar destabilising effect on mRIPK3 fibrils: hetero-amyloid assemblies are readily depolymerised by treatment with 2% SDS to yield monomeric mRIPK3 and M45 (**Fig. S17(c)**).

## Discussion

Since the first reports of functional amyloids, the participation of numerous partners has suggested the possibility that hetero-amyloids might play important roles in biology and dramatically widen the activities imparted by amyloids [31]. Examples of functional amyloid systems that depend on hetero-interactions include the fungal HET-S system for heterokaryon incompatibility, bacterial curli fibrils, and Drosophila cryptic RHIMs which function in the fruit fly immune system [32-35].These contrast with disease-associated amyloids, which are typically composed of a single type of polypeptide in each fibril [36].

The RIPK1:RIPK3 necrosome provided the first detailed study of a mammalian hetero-amyloid [4, 18], and both provided strong support for the hypothesis of hetero-amyloids serving as powerful signaling elements, and elucidated some the unique structural motifs that can be involved. It is proposed that the preference for the hetero-amyloid RIPK1:RIPK3 may account for the specificity in the signal transduction of necroptosis and that RIPK1:RIPK3 may seed the formation of homo-RIPK3 amyloids [18]. The structure of the minimum amyloid fibril core of human homo-RIPK3 was subsequently determined by cryo-EM and SSNMR [20]. In both cases, the RHIM-defining tetrad sequence of homo-RIPK3, VQVG, forms the essential central β strand of a small S-shaped fold. The homo structure has been proposed to be optimal for autophosphorylation of RIPK3 and activation of MLKL [20] based on its relatively small fibril pitch and large twist [37].

Viral RHIMs M45 from MCMV and ICP6 from HSV-1 have been shown to interfere with host necroptosis signaling [25, 38]; in addition, other viral RHIMs interact with host RHIMs to inhibit apoptotic pathways [39]. Evidence from in vitro experiments suggested that this activity was the result of the formation of hetero-amyloids containing host and viral RHIM proteins, in preference to those between host RHIM-containing proteins [25, 39]. The formation of hetero-amyloids depends significantly on the sequences of amyloid-forming proteins and in the present case, RIPK3, RIPK1, and M45 notably share conserved IQIG or VQVG motifs in their core tetrad regions that suggest the formation of β-sheet pairing in a cross species hetero-amyloid fold analogous to RIPK1:RIPK3. The structure of RIPK3(M45) is more similar to homo-RIPK3 than to RIPK3(RIPK1). Alignment of the hydrophobic cores (**Fig. 8(a)**) reveals that a segment of RIPK3 includes a remote RHIM-tetrad-like motif (sequence VNI) that is engaged with the core in the M45: RIPK3 and homo-RIPK3 presumably stabilizing the structure, while it is “released” from the amyloid core in RIPK1:RIPK3 possibly releasing it for interactions with other partners. Hydrophobic stabilizing interactions involving Ile and Val residues from M45 and RIPK3, respectively, and a Gln side chain ladder appear to also assist in the fibril formation leading to the feasibility of a M45:RIPK3 amyloid. The present study confirms such a proposed stacking. On the other hand, outside the core tetrad, several sequence mismatches occur in M45 relative to RIPK3 both upstream and downstream of the core region, disrupting interactions that are observed along the homo-amyloid fibril axis, such as asparagine or glutamine ladders and π-π stacking. Interestingly, we observed multiple arrangements of side chains in these sequence mismatches in the final ensemble of structures, as described in **Fig. 4(b)** and **(c)**, dynamic variation not seen in the previous homo- and hetero-amyloids, for example as seen in the altered peak intensity of Asn463 in homo-versus hetero-RIPK3. Direct measurements of dynamics, such as relaxation and order parameters, will be of interest to purse these issues quantitatively.

**Figure 8.**
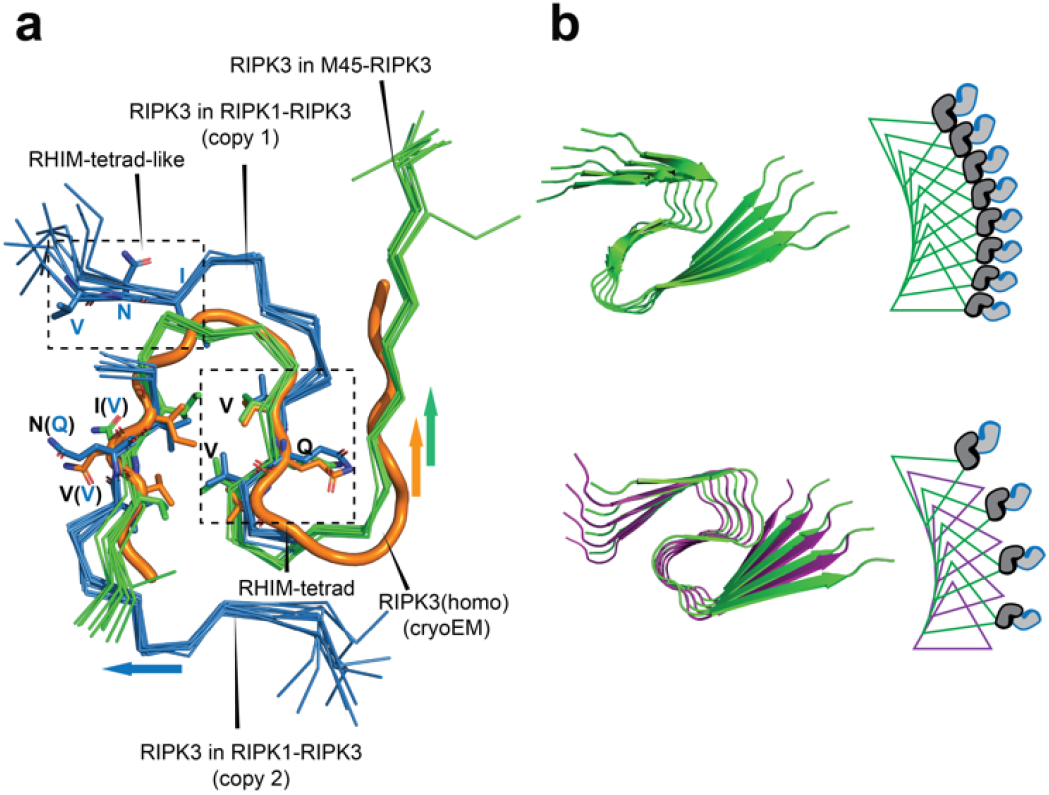
Hydrophobic cores of RIPK3 and schematic representation of the effect of interdigitation of M45 between RIPK3. (a) Alignment of RIPK3 structures in homo-RIPK3 (orange), RIPK1-RIPK3 (blue ribbons), and M45-RIPK3 (green ribbons). The residues used for alignment are V450-I452 and V458-V460 in homo-RIPK3 and M45-RIPK3. For RIPK1-RIPK3, two copies of the V458-V460 segment are utilized for alignment. Arrows indicate the sequence directions. (b) The position of RIPK3 kinase domains is determined by the pitch and twist of the amyloid fibril. The presence of the M45 RHIM in the heteromeric amyloid introduces a spacing of 4.8 Å between RIPK3 molecules and the interlayer twist angle of ∼9Å results in separation of RIPK3 kinase domains and prevents RIPK3 dimerization, autophosphorylation and subsequent MLKL activation.

Additional features suggest the presence of unanticipated stabilizing interactions that are unique to this hetero-amyloid. Charged residues play important roles providing specificity in the structure of this hetero-amyloid complex, and in general all amyloids. Frequently, in register homo-amyloid structures terminate at charged residues if there are no counterions to neutralize the net charge [40]. In other cases, charged residues face solvent, so that water molecules can attenuate this repulsion. This is observed in the structures of murine homo-RIPK3 determined by SSNMR, and human homo-RIPK3 fibrils characterized by cryo-EM, where E447 and D462 face the solvent [19, 20]. In the structure of human homo-RIPK3 determined by SSNMR, the sidechain of D462 appears to hydrogen bond to backbone amides [20]. In the hetero-amyloid studied here, the proteins exhibit high homology, allowing most hydrophobic residues to stack together and form hydrophobic cores. Additionally, some polar and neutral charged residues can stack together to form ladders involving Asn, Gln, Thr, and Ser, as demonstrated in the structures of RIPK1:RIPK3 and M45:RIPK3. In all the hetero-amyloids discovered so far, there are no same-charged residues colocalized [4, 25, 38]. In the alternating packing scheme, the distance between same-charged residues is doubled to approximately 9.6 Å. Meanwhile, the charged residues can interact with colocalized polar residues through hydrogen bonding, adding stability to the structure. In M45:RIPK3, the charged residues are further stabilized by a salt-bridge. It is noteworthy that the hydrophobic core formed by the β1 and β2 strands is sealed by this salt-bridge, a structural motif also observed in other amyloids, such as Aβ42 [41], as shown in **Fig. S18**. The stabilization offered by this interaction could compensate for the mismatches mentioned above or the potentially loosened interactions implied by NMR indications of increased dynamics. Murine RIPK3 does not have an aspartate residue in the position homologous to D462 (N452) but a glutamate residue is present immediately N-terminal of the murine RIPK3 core tetrad. The position of this charged residue makes the formation of a salt bridge with R53 of M45 unlikely and suggests that the mRIPK3 fold in M45:mRIPK3 hetero-amyloid may differ from that in homo-mRIPK3 [19]. Potentially the charged residues for the murine case also somehow provide energetic preference for the hetero-amyloid, and this subject would be interesting for further investigation.

Our SSNMR analysis and mutational data provide a possible explanation of why M45 can disrupt the formation of the human RIPK1-RIPK3 necrosome. Firstly, the anti-necroptotic M45:RIPK3 hetero-amyloid forms preferentially as compared with RIPK1:RIPK3. This may be in part because larger regions of M45 and RIPK3 adopt β sheet structure in the human:viral RHIM hetero-amyloid than in RIPK1:RIPK3. Furthermore, the formation of the salt bridge in the hetero-amyloid may result in a higher preference of RIPK3 forming an amyloid with M45 than with RIPK1. The mutational data provides evidence for this more extensive interface between RIPK3 and the viral protein. Mutations across M45 spanning from R53 to A72 disrupt the ability of the viral protein to form a hetero-amyloid with RIPK3. Meanwhile, in the pro-necroptotic RIPK1:RIPK3 complex, significant regions are released from the amyloid core that might engage in unspecified downstream interactions.

Steric factors may also play a role in the inhibitory M45:RIPK3 complex. The activity of M45 prevents autophosphorylation of RIPK3 and its subsequent phosphorylation of MLKL [25, 42]. The interdigitation of M45 between RIPK3 molecules in the hetero-amyloid will have a marked effect on the relative positioning of RIPK3 molecules (**Fig. 8(b)**). RIPK3 molecules in the hetero amyloid will be ∼9.6 Å apart instead of 4.8 Å and the twist along the fibril will impose an additional change in the relative orientation of the kinase domains. The M45:RIPK3 structure determined here by SSNMR exhibits an average twist of ∼9° between alternating layers. The increased separation of kinase domains and the presence of the large ribonucleotide reductase domain from M45 may attenuate the ability of RIPK3 to autophosphorylate or activate MLKL.

In conclusion, this study elucidates the inhibition of necroptosis by M45 by demonstrating its ability to form a hetero amyloid with RIPK3, thus competing with the other pro-necroptotic amyloids that RIPK3 forms. Notably, the complex of M45 and RIPK3 is novel in that it involves proteins from two species, pathogen and host, and its formation serves to allow the pathogen to evade the host immune response.

## Materials and Methods

### Expression and Purification of Unlabeled and Isotopically Labeled M45 and RIPK3

RHIM-containing portions of MCMV M45 (Q06A28; residues 1–90) and human RIPK3 (Q9Y572; residues 387–518) were produced as His_6_-ubiquitin-RHIM fusion proteins using the pHUE vector system [43], i.e. His_6_-Ub-M45_1–90_ and His_6_-Ub-RIPK3_387–518_. Both constructs have a 9-residue linker with a TEV site between ubiquitin and the RHIM-containing portion and their amyloid-forming properties have been reported previously [25]. Successful cloning was confirmed by sequencing at the Australian Genome Research Facility at the Westmead Millennium Institute. Proteins were expressed in BL21(DE3) *E. coli* grown in minimal media with natural abundance (n.a.) or _15_N-ammonium chloride and n.a. or U-^13^C, 1,3-^13^C or 2-^13^C glycerol to produce natural abundance or partially isotopically labeled samples.

Proteins were expressed in 1 L batch cultures according to the high cell density protocol [44]. The process followed 300 μL freshly transformed *E*.*coli* BL21*(DE3) inoculated into 10 mL ModC1 minimal medium, incubated overnight and cell suspension diluted 5x in fresh ModC1 medium (40 g/L n.a. glycerol or 20 g/L glycerol-^13^C_3_ 99 atom % ^13^C or 5-10 g/L glycerol-1,3-^13^C2 99 atom% ^13^C or 5-10 g/L glycerol-2-^13^C 99 atom% ^13^C, 5.16 g/L n.a. NH_4_Cl or ^15^NH_4_Cl ≥98 atom% ^15^N) and grown at 37°C for two-three OD_600_ doublings. The cells were inoculated into fresh n.a. or labeled-ModC1 to a volume of 100 ml and grown to OD_600_ of between 0.4-1.9 dependent on growth medium before inoculation into 900 ml labelled expression medium in a bioreactor. All cultures were induced for expression by the addition of IPTG to a final concentration of 1 mM, with induction at 20°C. Dependent on the labelling scheme further NH_4_Cl was added to the culture. Cell suspensions were pelleted by centrifugation at 8000g, 4°C for 20 min and biomass stored at −80 °C.

For purification of His_6_-Ub-M45_1–90_ and His_6_-Ub-RIPK3_387–518_ proteins, cells were lysed by sonication in 20 mM Tris.HCl, 1 mM EDTA, pH 8.0 and the insoluble material pelleted by centrifugation. The pellets were resuspended in 20 mM Tris.HCl, 6 M GuHCl, pH 8.0, and the mixture was sonicated and then stirred for 3 h at RT before storage at 4 °C overnight. Soluble material was collected by centrifugation at 20,000*g* for 1 h and His-tagged proteins were purified using Ni-NTA affinity chromatography (Life Technologies), with exchange into 100 mM NaH_2_PO_4_, 20 mM Tris.HCl, 8 M urea, and 5 mM β-mercaptoethanol at pH 6.5 for washing and at pH 4.0 for elution from the Ni-NTA agarose.

To generate M45 and RIPK3 homomeric fibrils or M45-RIPK3 isotopically labeled hetero-amyloid, His-Ub-RIPK3_387–518_ and M45_1–90_ alone or mixed together in 1:1 molar ratios at 50 µM or with an appropriate volume of 8 M urea-containing buffer, were dialysed for 1 hour at room temperature against a buffer comprising 25 mM NaH_2_PO_4_, 150 mM NaCl, 0.5 mM DTT, pH 7.4. At this point, TEV protease was added, and dialysis continued overnight at room temperature. SDS-PAGE was performed to confirm cleavage of the affinity tag and this demonstrated that during dialysis and incubation with TEV, the RHIM-containing fragments self-assembled into insoluble material (Fig. S1). Following centrifugation, the supernatant, containing the cleaved His_6_-ubiquitin, was separated from the pellet. The pellet, containing the RHIM fragment(s), was washed three times with MilliQ water and then snap-frozen in liquid nitrogen and lyophilized. The lyophilized fibril samples were resuspended in 20 mM HEPES, 20 mM NaCl, 1 mM TCEP.HCl, pH 7.4 and then sonicated with a probe sonicator, on ice, with three 20-second bursts, to resuspend and hydrate the fibrils. Sodium azide was added to a final concentration of 0.1%.

Protein samples generated in this way: n.a. M45; n.a. RIPK3; ^15^N-M45; ^15^N-RIPK3; U-^13^C,^15^N-M45; 1,3-^13^C,^15^N-M45; 2-^13^C,^15^N-M45; U-^13^C,^15^N-RIPK3; 1,3-^13^C,^15^N-RIPK3; 2-^13^C,^15^N-RIPK3. In all of these samples, M45_1–90_ and RIPK3_387–518_ are indicated as M45 or RIPK3, for clarity.

### Expression and purification of YPet and mCherry-tagged M45 and RIPK3 constructs

Synthetic genes encoding WT and AAAA mutant forms of M45_1-90_, human hRIPK3_387–518_ and murine mRIPK3_379-486_ regions, and YPet and mCherry, were purchased from Genscript and cloned into a modified pET15b vector to generate vectors encoding His_6_-YPet-hRIPK3_387–518_, His_6_-YPet-mRIPK3_379-486_ or His_6_-M45_1–90_-mCherry. Polyhistidine-tagged fluorescent protein-RHIM fragment fusion proteins were expressed and purified as described in [25]. These constructs are referred to as YPet-hRIPK3, YPet-mRIPK3 and mCherry-M45 for clarity.

### Solid State NMR Experiments and Data Processing

Fibrillar samples for solid-state NMR assignments were produced using a partial isotopic labeling scheme wherein one protein is uniformly labeled with ^13^C and ^15^N while the other protein is at natural abundance (n.a.), e.g., U-^13^C,^15^N-M45:n.a. RIPK3 and vice versa. The NMR resonance assignments were conducted using two and three-dimensional (2D and 3D) correlation experiments, including homonuclear carbon-carbon experiments such as DARR [45] and CORD [46], as well as the traditional backbone walk experiments: NCOCX, NCACX, and CANCO [47]. A detailed summary of the experiments can be found in Supplementary **Table S9** and **S10**. Analyzing these datasets, we determined unambiguous backbone and sidechain chemical shifts of the observed residues, which are summarized in **Fig. S4** and **Table S7** and **S8**.

We used an alternately labeled sample, as above, using 1,3-^13^C-glycerol and 2-^13^C-glycerol labeling schemes [48] on the isotopically labeled protein to observe intramolecular medium and long-range contacts. Using both carbon homonuclear (DARR) and C-N heteronuclear (TEDOR) [49] experiments (detailed in **Table S9** and **S10**). The contacts are summarized in **Table S2** and two representative DARR spectra are shown in **Fig. S5 (a)** and **(b)**.

For the examination of intermolecular contacts in the hetero-amyloid samples were made by mixing 1,3-^13^C-glycerol and 2-^13^C-glycerol labeled proteins, e.g., 1,3-^13^C-glycerol-M45:2-^13^C-glycerol-RIPK3 and referred to herein as 1,3-M45:2-RIPK3. These samples should lead to additional NMR resonances that are due to intermolecular contacts within the complex.

MAS spectra were acquired using a Bruker Avance NEO 17.6 T (750MHz ^1^H frequency) spectrometer equipped with a 1.9mm HXY probe and a Bruker Avance NEO 21.1 T (900MHz ^1^H frequency) spectrometer with a 3.2mm E-free or 1.6mm E-free probe. All experiments were conducted at approximately 280 K with 100 kHz SWf-TPPM [50] ^1^H decoupling unless specified otherwise. A detailed summary of experimental conditions and processing parameters can be found in the supplements **Table S9** and **S10**.

All NMR data were processed using Topspin. NMR chemical shift assignment and analysis were conducted using CCPNMR version 2 [51].

### Structural Calculations of M45-RIPK3

Structure calculations were carried out using simulated annealing with Xplor-NIH. 4 copies of M45 and 4 copies of RIPK3 were included into the calculations. M45_48-75_ and RIPK3_447-473_ were included in the calculations since these residues are assigned. Non-crystal symmetry was applied on M45 and RIPK3 respectively, since we only observed one set of chemical shift values for M45 and RIPK3(except I452). Inter-strand C-C distances of 4.75 ± 0.1 Å were enforced using distance constraints for the residues showing high probabilities of *β*-sheet in both M45 and RIPK3. These included residues 51-73 in M45 with the exception of V58, G60, and H66, and residues 448-470 in RIPK3 excluding C455, G457, and N463. Backbone dihedral angle constraints are based on predictions from both TALOS+ [52] and TALOS-N [53]. Only dihedral predictions within a 30-degree difference are included in the calculations, and the constraints used in the calculation encompass the full range of predictions from both methods.

Two rounds of annealing were performed. In the initial round, high-temperature dynamics were initiated at 4000K for 100ps or 1000 steps in the torsion angle space. Subsequently, annealing occurred to reach 25K, with a temperature decrement of 12.5K, and at each temperature, dynamics in the torsion angle space were run for 2ps or 1000 steps. Ultimately, 500 steps of energy minimization were conducted in both torsion angle and Cartesian coordinates. In the first round of annealing, unambiguous intramolecular and intermolecular contacts are employed as distance-based constraints, namely NOEs. Contacts derived from DARR have an upper limit of 7.5 Å, and contacts derived from TEDOR have an upper limit of 6.5 Å. Sequential contacts were also included, as summarized in **Table S4**. In terms of unambiguous intermolecular contact, they are encoded as ambiguous contacts in a way that one atom from M45 can be in contact with the other atom from RIPK3 either below or above the M45 layer. During the initial annealing phase, the energy function incorporates essential Xplor terms including BOND, ANGL, and IMPR, which account for covalent bond lengths, bond angles, and improper dihedral angles, respectively. Additionally, knowledge-based energy terms encompassing hydrogen bond interactions, and a torsion angle database are also integrated. Moreover, a purely repulsive nonbonded term is also incorporated into the energy function. Hydrogen bonds between R53-HN and D462-O*δ* are enforced given the observation of the cross peak between R53-C*ζ* and D462-C*γ*, along with the chemical shift variations observed in D462-C*γ* when compared with homo-RIPK3.

In the first approach for assigning intermolecular contacts (as indicated by blue circles in **Fig. 3 scheme I**), M45 was considered unambiguous for the initial folding under Scheme I. To distinguish the BA or BC stacking preferences for each unambiguous intermolecular contact, we computed the difference in the distance between the nuclei in both BA and BC for ten unambiguous intermolecular cross-peaks within the 160 lowest-energy structures, selected from the pool of 320 calculated structures during the initial folding process (**Fig. S10(A)**). We observed that the R53 (M45)-D462 (RIPK3) contacts had two distinct distributions. The majority distribution was centered around −6 Å, representing structures where the contact was between segments B and A, while the minority distribution, with contacts between segments B and C, was centered near +5.5 Å. Consequently, we classified these 160 structures into two ensembles based on this contact: the major BA species (around 75%) and the minor BC species (around 25%).

We found two additional contacts that also displayed a preference between the BA and BC assemblies, albeit weaker than that of the R53Cζ(M45)-D462Cγ(RIPK3) contact: V58Cγ1(M45)-C458Cβ(RIPK3) and I61Cβ(M45)-C455Cβ(RIPK3). The former exhibited a preference similar to that of the R53Cζ(M45)-D462Cγ(RIPK3) contact, while the latter exhibited the opposite preference.

We further examined the backbone RMSD alignment among these 160 structures with respect to the lowest energy structure, as depicted in **Fig. S10 (b)** and **(c)**. The major species (BA contact) exhibited an average backbone heavy atom RMSD of 1.3 Å and a similar energy, while the minor species showed an average RMSD of 3 Å. The alignment was performed between residues 50-72 on M45 and 447-469 on RIPK3. When we ran the calculation using the second approach, where RIPK3 was considered unambiguous (as indicated by orange circles in **Fig. 3 scheme I**), the results were remarkably similar, as shown in **Fig. S12**. The minor species accounted for approximately 25% of the structures using both approaches.

For assigning intermolecular contacts, we also considered another approach where we imposed additional ambiguous constraints on the flanking segment. However, in this approach, a segment would be constrained twice, which introduces another artifact.

Following the calculation of 320 independent structures in the initial round, the 10 lowest-energy structures, including one from the minor species, were retained for subsequent refinement. Using the 10 best structures from the initial folding and our chemical shift assignments, we predicted the M45-RIPK3 intermolecular contacts expected in the long mixing DARR spectra, applying a distance threshold of 7 Å. By comparing these predictions with the experimental spectra (**Figs. S6– S8**), we identified an additional 6 unambiguous M45-RIPK3 intermolecular contacts, which we subsequently included in the refinement. In the second round of annealing, most settings remained consistent with those used in the first round. Additionally, low-ambiguity contacts derived from DARRs were incorporated as ambiguous distance constraints. Residue affinity potential was also included to account for the arrangement of hydrophobic, polar, and charged residues. Each of the 10 initial structures generated an additional 10 independent structures, resulting in a total of 100 structures.

Improved structural convergence was demonstrated in this second round of annealing, as shown in **Fig. S13**, where among the 90 lowest-energy structures without violations, 95% were classified as major species. Further minimization was subsequently performed on the 10 lowest-energy structures. These 10 structures underwent an all-atom minimization process comprising 500 steps, with constraints applied to maintain the backbone RMSD within a 1 Å threshold.

In Schemes II and III, inter-strand C-C distances of 4.75 ± 0.1 Å are enforced between neighboring segments in homo-M45 and homo-RIPK3. In Scheme II, intermolecular constraints are applied only at the interface between homo-M45 and homo-RIPK3 along the fibril axis, whereas in Scheme III, they are applied at every layer of M45 and RIPK3. A total of 320 structures are calculated in the initial folding using the same protocol described before.

### Cryo-EM of M45 homomeric fibrils

M45 protein was purified for cryo-EM as for ssNMR but after fibrils were formed by dialysis and His-Ub removed by TEV treatment, the fibrils were pelleted by centrifugation, washed with water, and treated with 100 % formic acid to dissociate assembled M45 back to a monomeric state. Formic acid was removed by lyophilisation and the protein was further purified by RP-HPLC, snap-frozen in liquid nitrogen, then lyophilised and stored at −20 °C until required. The M45 protein was resolubilised in 1 mM Acetic acid, 0.2 % TFA, 1 mM TCEP, pH 1.5 to a final concentration of 200 µM and then dialysed against 38.7 mM Acetic acid, 11.3 mM Sodium Acetate, pH 4.0 overnight at room temperature, before being diluted 10-fold in fresh buffer.

Samples of diluted M45 fibrils were then applied to AuFlat GF-1.2/1.3-3Au-45 nm grids treated with a positive polarity glow discharge and vitrified in liquid ethane using a Vitrobot Mark IV (ThermoFisher Scientific). Micrographs were acquired using a 300 keV Titan Krios microscope (ThermoFisher Scientific) equipped with an energy filtered BioQuantum^®^ K3 camera (Gatan) with a slit width of 10 eV. Further details are provided in **Table S12**.

### Helical reconstruction

Movie frames were gain-corrected, aligned, dose-weighted and summed using the motion correction program implemented in RELION-4.0 [54]. The contrast transfer function (CTF) was then estimated from the motion-corrected micrographs using CTFFIND-4.1 [55]. Further image-processing and helical reconstruction was performed using methods in RELION-5.0 [56, 57]. Isolated fibrils of M45 were picked manually from 14,767 micrographs and subjected to several iterations of reference-free 2D classification to remove suboptimal fibril segments. An initial 3D model was then constructed by producing a sinogram from selected 2D class averages which collectively span a single crossover of the helical twist, as previously described [58]. Numerous rounds of masked 3D autorefinement and 3D classification were then performed to further remove suboptimal fibril segments and optimise helical parameters. These refinements were also performed using Blush regularisation [59] where beneficial to reduce noise and improve map resolution and was particularly useful in preventing high noise arising from overfitting in conventional refinement methods. Iterative Bayesian polishing and CTF refinement was also performed, as previously described, with 3D autorefinement and, if beneficial, 3D classification repeated. Final maps were sharpened using the post-processing method implemented in RELION-5.0 with resolutions estimated from the resulting Fourier shell correlations of 0.143 between independently refined half-maps, as described previously [60]. Further details are provided in **Table S12**.

### Thioflavin T fluorescence assays of amyloid assembly

Protein from stock solutions containing 8 M urea was diluted to 2.5 μM in 25 mM NaH_2_PO_4_, 150 mM NaCl, 40 μM ThT, 0.5 mM DTT, pH 7.4 to a final volume of 200 μL in a Costar 96-well fluorescence plate (Corning #3631). The residual urea concentration was less than 300 mM. Immediately after dilution, the plate was transferred to a POLARstar Omega microplate reader (BMG Labtech) maintained at 37 °C and samples were mixed by shaking at 500 rpm for 30 seconds, then fluorescence intensity (Ex/Em 440/480 nm) was recorded every 60 seconds for 2.5 h. Recorded data were analyzed in Microsoft Excel and GraphPad Prism.

### Negative stain transmission electron microscopy

Fibril samples for transmission electron microscopy were prepared in 25 mM phosphate, 150 mM NaCl, pH 7.4. 4 (2.1.2). Formvar- and carbon-coated copper grids (200 mesh, ProScitech) were floated on the protein droplets for 30 seconds, then excess solution was removed with filter paper. Grids were washed three times with filtered water and then stained with 2% uranyl acetate before air-drying and imaging in a Technai T12 electron microscope operating at 120 kV fitted with an Olympus Veleta CCD Camera.

### Two-Colour Coincidence Detection spectroscopy

Homomeric or heteromeric samples of fluorescently tagged NYPet-RIPK3 and M45-mCherry constructs were prepared at 35 μM in 25 mM phosphate, 150 mM NaCl, 8 M urea, 0.5 mM DTT, pH 7.4 and then diluted 100-fold to 0.35 µM with 25 mM phosphate, 150 mM NaCl, 0.5 mM DTT, pH 7.4 and immediately examined in the confocal microscope. The fluorescence emission of both fluorophores was recorded simultaneously over a period of 3 minutes. Data were analyzed and plotted in GraphPad Prism or Excel. Interaction between RIPK3 (WT) and M45 (WT and mutants) was characterized by Two-Color Coincidence Detection (TCCD). The ratio R is calculated as:

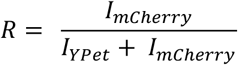

Where *I*_*mCherry*_ and *I*_*YPet*_ are the intensities collected from individual bursts after corrections for direct excitation of mCherry (2%) and leakage of the YPet into the mCherry detection channel (6%). Bursts were identified when the signal (*I*_*mCherry*_ + *I*_*YPet*_) is >3000 photons per ms above background. A ratio between 0.15 and 0.5 indicates coincident detection. The coincidence is therefore calculated as the total percentage of events with 0.15<R<0.5.

### SDS agarose gel electrophoresis

Single proteins (5 µM) or mixtures of proteins (RIPK3:M45, 5:20 µM) were prepared in 25 mM phosphate, 150 mM NaCl, 8 M urea, 0.5 mM DTT, pH 7.4 and dialysed overnight to allow assembly of fibrils. Fibrils and monomeric controls were prepared with 4 % glycerol and 0.0008 % bromophenol blue and incubated with either 0 % or 2 % SDS for 10 minutes at room temperature. Samples were electrophoresed through a 1 % agarose gel in TAE buffer containing 0.1 % SDS for 2 hours at 80 V. Gels were imaged with a Bio-Rad ChemiDoc Imaging System using 605/50 and 695/55 nm emission filters, corresponding to the emission wavelengths of YPet and mCherry and images were processed in ImageJ.

## Acknowledgments

The work was funded by the National Science Foundation (NSF MCB 24-27038) and by the Australian Research Council Discovery Project (DP DP200102463) and NHMRC Ideas Grant (APP2019231). The NMR data were collected at the New York Structural Biology Center (NYSBC) with support from by NIH grants S10OD030373 (900 MHz) and P41GM118302 (750 MHz). A.E.M. is a member of the NYSBC. We thank Dr. Charles D. Schwieters from NIDDK for his help with structure calculation using Xplor-NIH. We acknowledge the support of the Australian Government in provision of access to ANSTO’s National Deuteration Facility, which is partly funded through the National Collaborative Research Infrastructure Strategy (NCRIS), for production of isotopically labelled proteins via NDF proposals NDF7209 and NDF8102. NRV was supported by a Research Training Program Scholarship from the Australian Government (Department of Education, Skills and Employment). The authors acknowledge the facilities and the assistance of staff within the Sydney Analytical and Sydney Microscopy & Microanalysis Core Research Facilities at the University of Sydney. The authors thank Dr. Benjamin Ryskeldi-Falcon and the Scientific Computing department at the MRC Laboratory of Molecular Biology for access to the computing resources required for cryo-EM data processing.

## Author Contributions

Chengming He: Methodology, study design, investigation, formal analysis, visualization, writing – original draft, review, and editing.

Nikhil R. Varghese: Methodology, study design, investigation, formal analysis, visualization, writing – review and editing.

Eric G. Keeler: Methodology, study design, investigation, writing – review and editing. Chi L. L. Pham: Methodology, study design, investigation, writing – review and editing.

Brayden Williams: Methodology, study design, formal analysis, visualization.

Stephan Tetter: Investigation, writing – review and editing.

Crystal Semaan: Investigation, resources.

Karyn L. Wilde: Resources, writing – review and editing.

Simon H. J. Brown: Investigation, resources, writing – review and editing.

James C. Bouwer: Investigation, resources, writing – review and editing.

Yann Gambin: Resources, formal analysis, visualization, writing – review and editing.

Emma Sierecki: Resources, formal analysis, visualization.

Megan Steain: Conceptualization, writing – review and editing, funding acquisition.

Margaret Sunde: Conceptualization, methodology, study design, writing – original draft, review and editing, visualization, project administration, supervision, funding acquisition.

Ann E. McDermott: Conceptualization, methodology, study design, writing – original draft, review and editing, visualization, project administration, supervision, funding acquisition.

