## Supplementary data for "Structural Studies of an Anti-necroptosis Viral:Human Functional Hetero-amyloid M45:RIPK3 using SSNMR"

**Supplementary information**

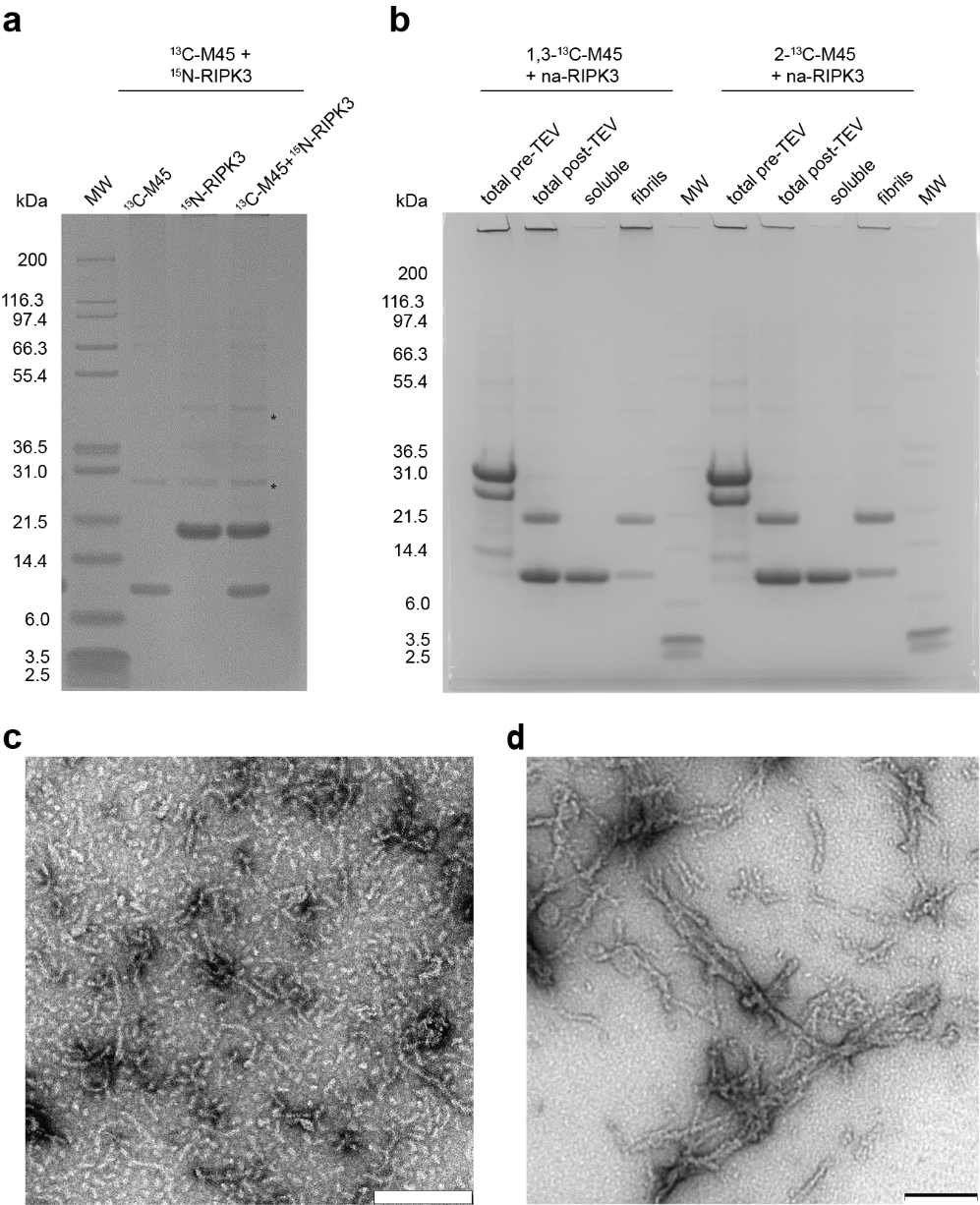

**Figure S1.** Biophysical characterization of NMR and fibril samples. (a) SDS PAGE analysis of fibril samples prepared from ^13^C-M45, ^15^N-RIPK3 or a 1:1 mixture of the two proteins, following removal of His-Ub tag. Asterisk indicates two bands arising from n.a. TEV. (b) Preparation of heteromeric fibrils containing 1,3-^13^C-M45 and n.a.-RIPK3, or 2-^13^C-M45 and n.a.-RIPK3. SDS PAGE analysis of soluble His-Ub-M45 and His-Ub-RIPK3 protein mixture before and after cleavage with TEV protease, followed by separation of supernatant and insoluble, pellet fractions. (c) Transmission electron micrograph of heteromeric fibrils prepared from ^13^C-M45 and ^15^N-RIPK3 following removal of His-Ub tags. (d) Transmission electron micrograph of heteromeric fibrils prepared from His-YPet-RIPK3 and His-M45-mCherry.Scale bar 200 nm in both TEM images.

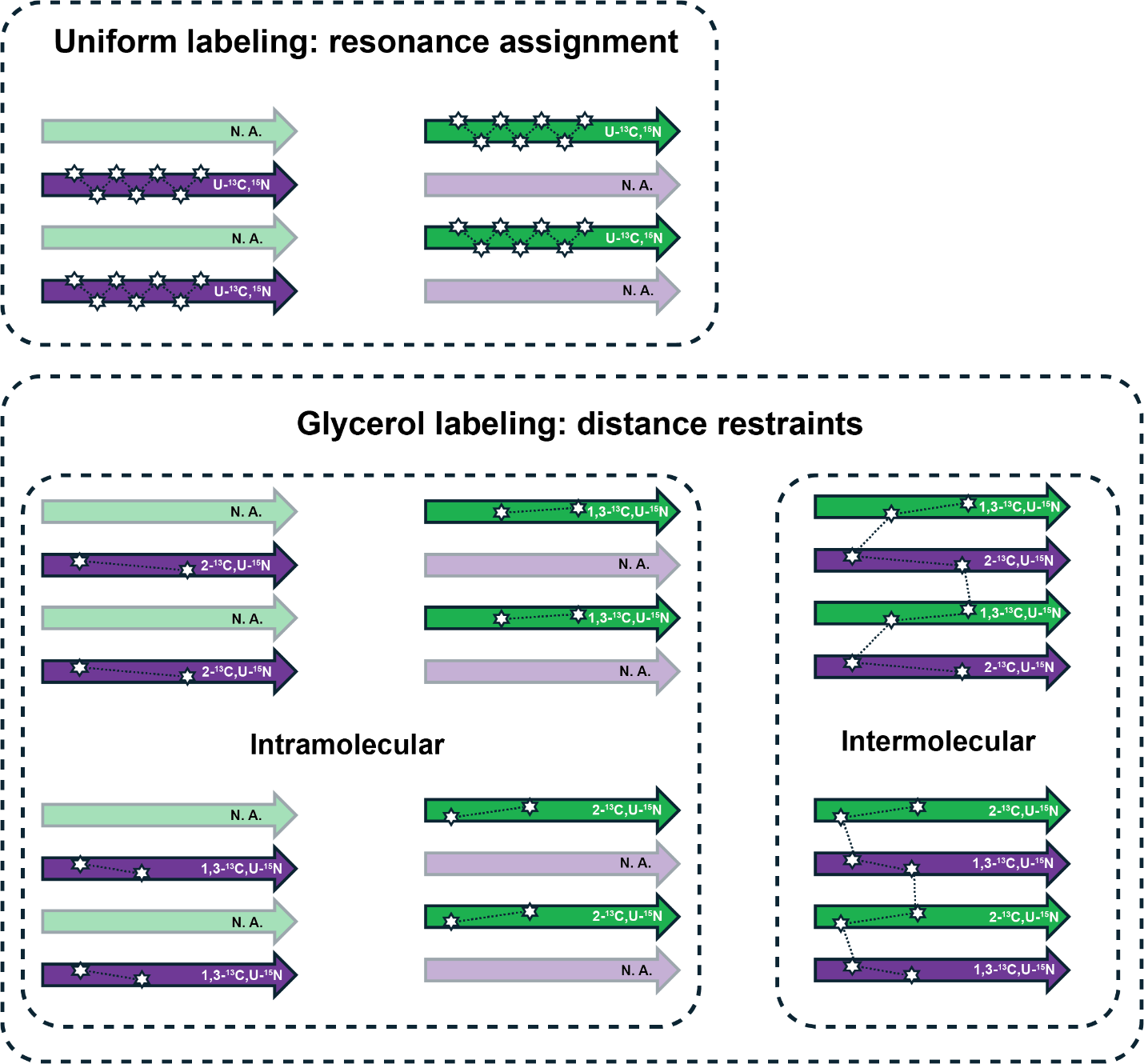

**Figure S2.** Illustration of labeling schemes. Solid colors represent isotopic labeling, while faded colors indicate natural abundance (N.A.). Uniform labeling is used to assign resonances in M45 and RIPK3. Glycerol labeling is employed to identify distance restraints. Single-labeled samples are used for intramolecular contacts, and doubly mixed-labeled samples are used for intermolecular contacts. Each labeling scheme is designed to detect a specific set of contacts, as indicated by the black dotted lines.

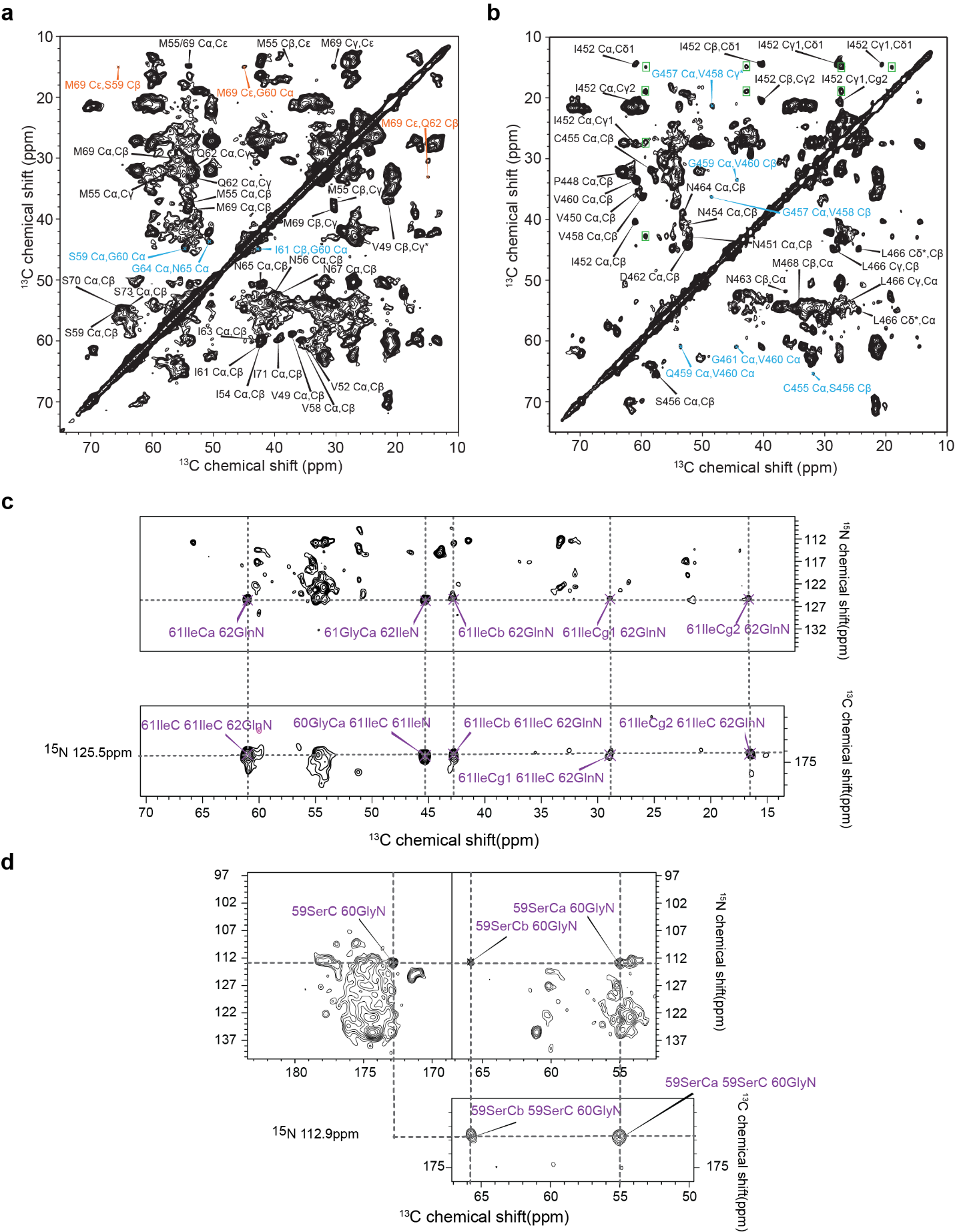

**Figure S3.** DARR spectra and 3D slices of uniformly labeled samples. (a) 75ms DARR spectrum of U-^13^C,^15^N-M45 / n.a. RIPK3, where intra-residue, sequential, and long-range cross peaks are colored in black, blue, and orange, respectively. (b) 75ms DARR spectrum of n.a.M45 / U-^13^C,^15^N-RIPK3, using the same color scheme as (a). (c) 2D N(CO)CX and a slice of NCOCX at ^15^N 125.5 ppm of U-^13^C,^15^N-M45, indicating the backbone and sidechain correlations from G60 to Q62. (d) Same experiments as (c) with ^15^N 112.9 ppm showing the correlations from S59 to G60.

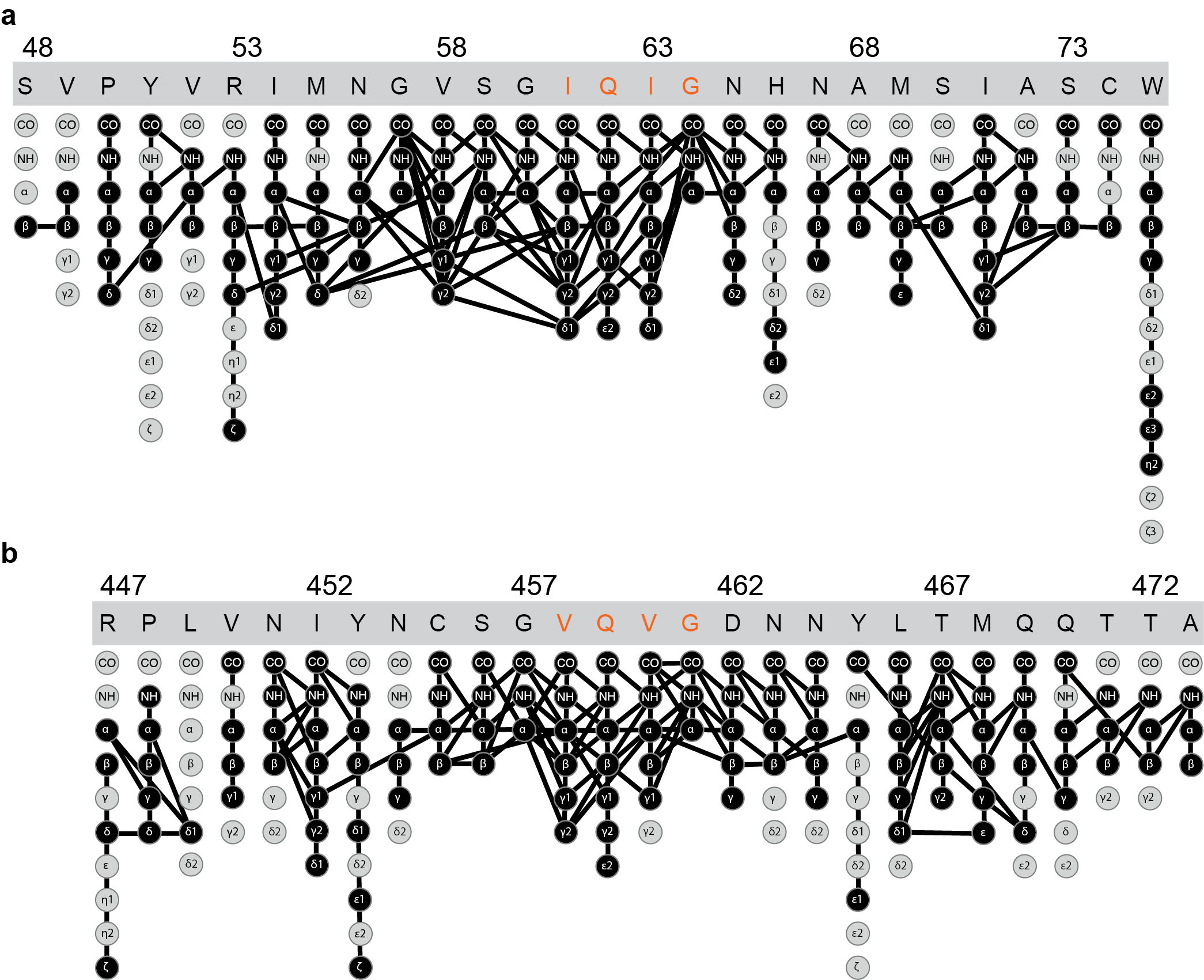

**Figure S4.** Assignment graph of M45 and RIPK3. (a) and (b) represent assignment graphs of M45 and RIPK3, respectively, where assigned atoms are labeled in black. Contacts within 4 residues are indicated by black lines.

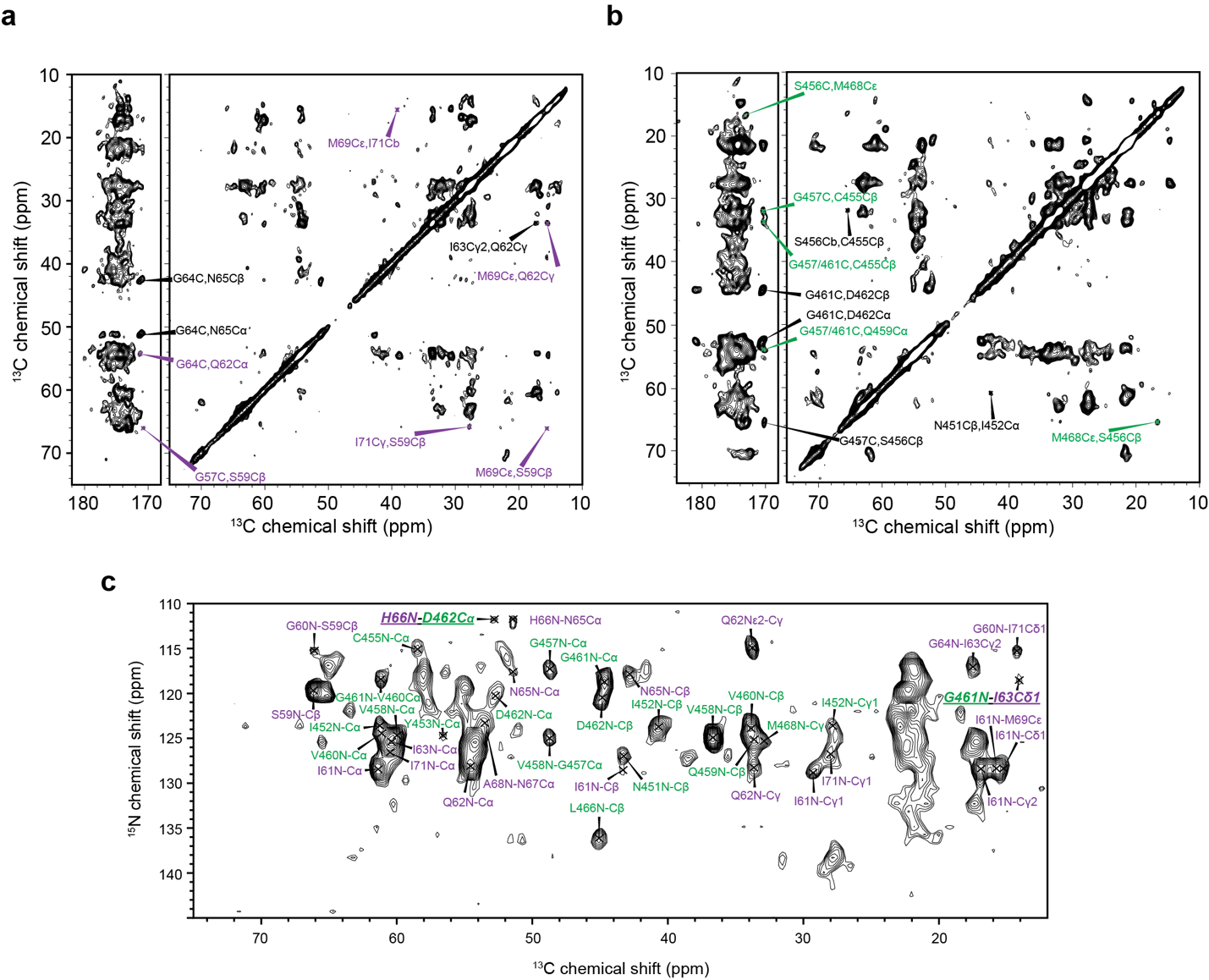

**Figure S5.** Spectra of glycerol labeled samples. (a) 350ms DARR of 1,3-^13^C-glycerol,^15^N-M45 : n.a. RIPK3, where sequential contacts are labeled in black and medium/long-range contacts are labeled in purple, respectively. (b) 350ms DARR of n.a. M45 : 1,3-^13^C-glycerol,^15^N-RIPK3, where sequential contacts are labeled in black and medium/long-range contacts are labeled in green, respectively. (c) 7.2ms TEDOR of 1,3-^13^C-glycerol,^15^N-M45 : 2-^13^C-glycerol,^15^N-RIPK3. M45 and RIPK3 intramolecular contacts are labeled in purple and green respectively. M45-RIPK3 intermolecular contacts are bold and underlined.

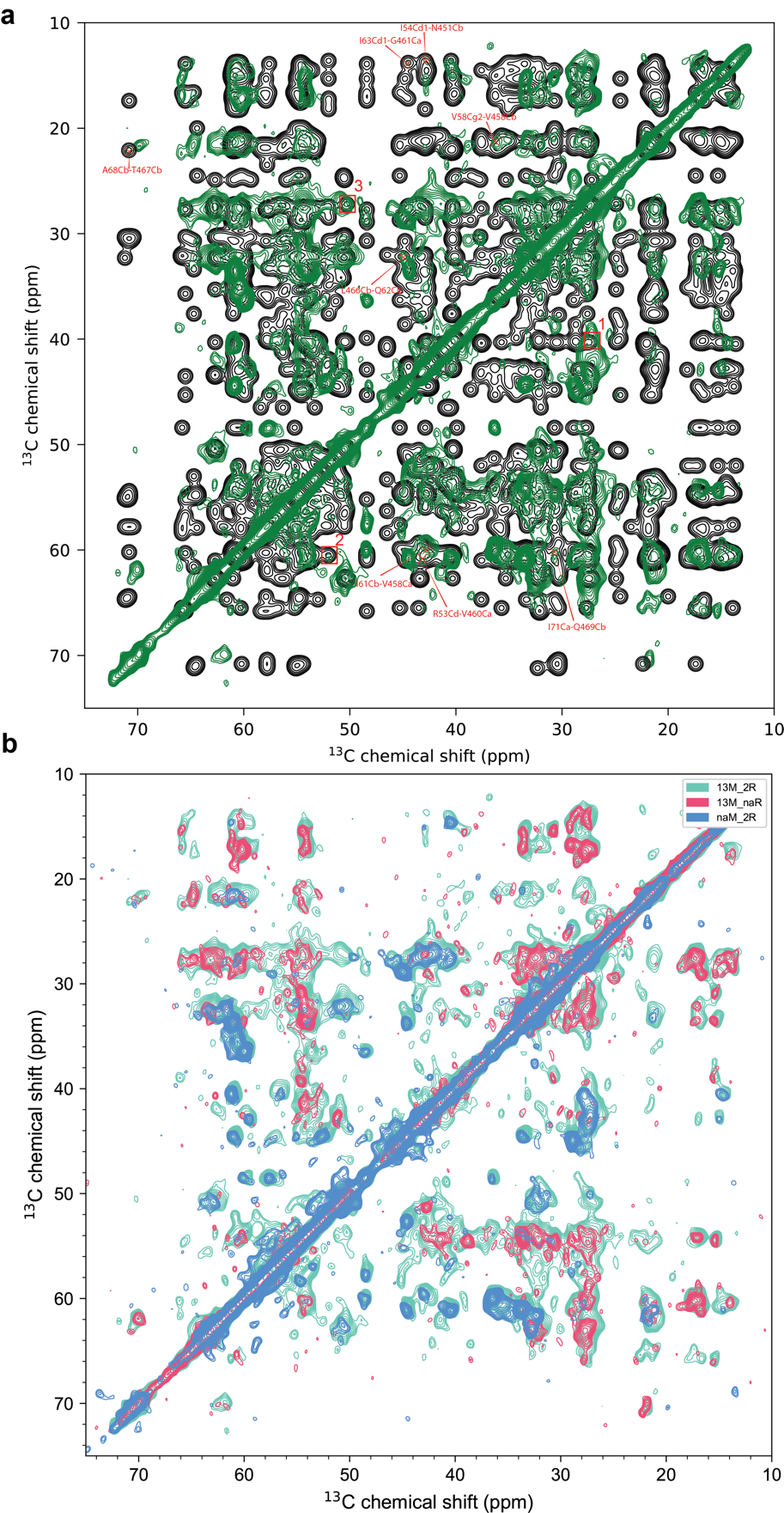

**Figure S6. Simulated and experimental spectra of 1,3-M45:2-RIPK3.** (a) Simulated spectra (black) and 500 ms DARR spectra of 1,3-¹³C-M45:2-¹³C-RIPK3 (green). (b) Overlay of 500 ms DARR spectra of 1,3-M45:2-RIPK3 (green), 1,3-M45:na.-RIPK3 (red), and na.-M45:2-RIPK3 (blue). The first contour is plotted as 3 times of noise.

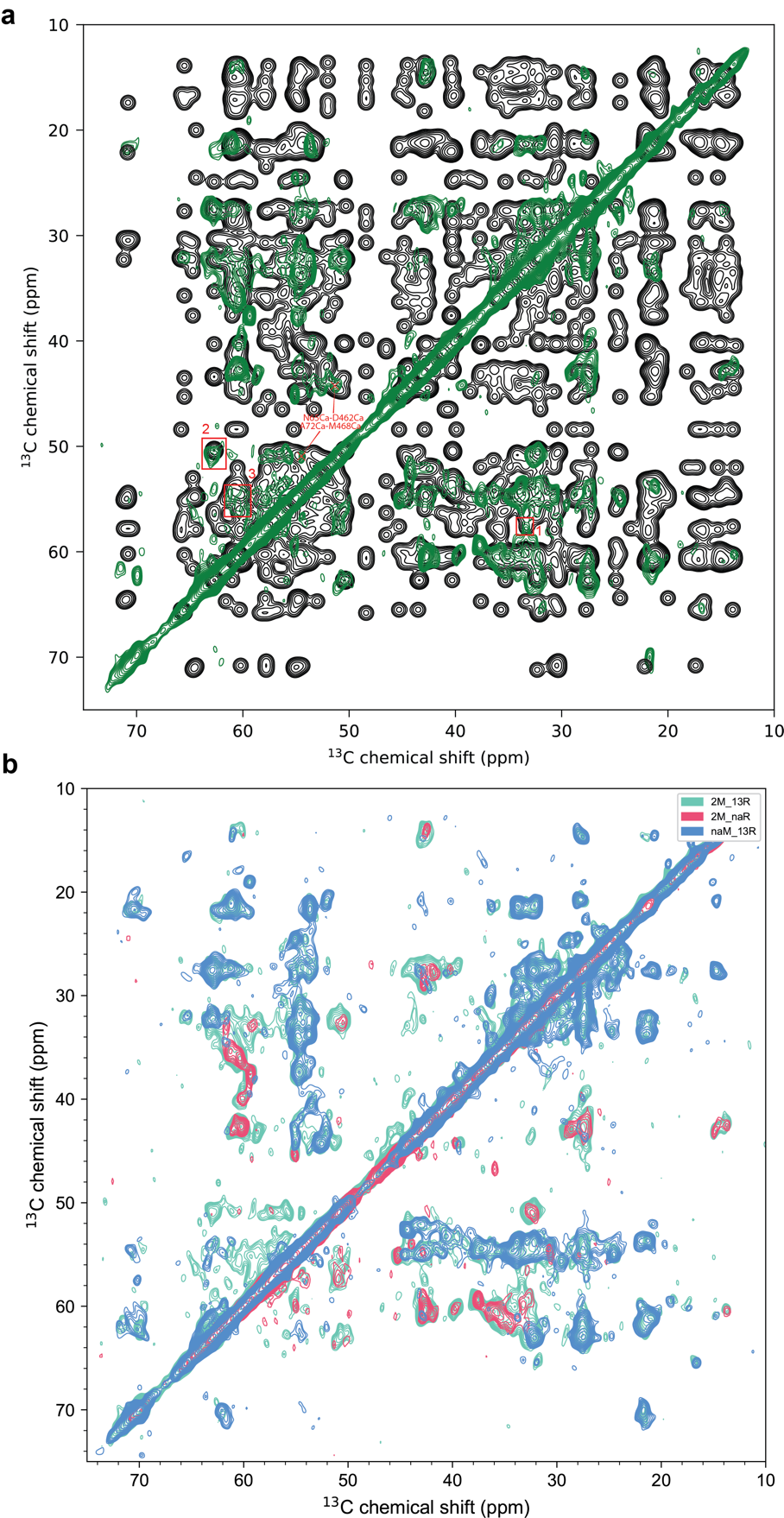

**Figure S7. Simulated and experimental spectra of 2-M45:1,3-RIPK3.** (a) Simulated spectra (black) and 350 ms DARR spectra of 2-¹³C-M45:1,3-¹³C-RIPK3 (green). (b) Overlay of 350 ms DARR spectra of 2-M45:1,3-RIPK3 (green), 2-M45:na.-RIPK3 (red), and na.-M45:1,3-RIPK3 (blue). The first contour is plotted as 3 times of noise.

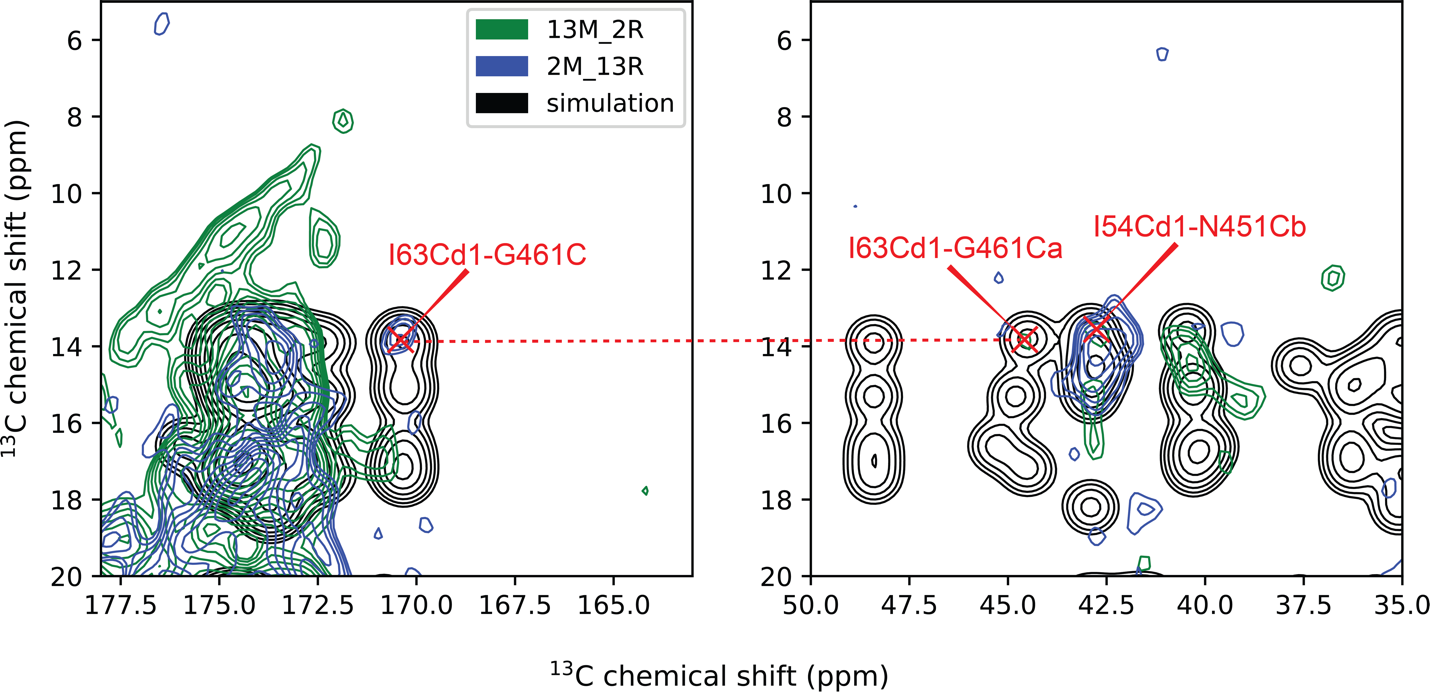

**Figure S8. Cross-validation of weaker intermolecular peaks.** Overlay of DARR spectra of 1,3-M45:2-RIPK3 (green), 2-M45:1,3-RIPK3 (blue), and simulated spectra (black). The highlighted peaks in the blue spectra support the weaker signals observed in the green spectra.

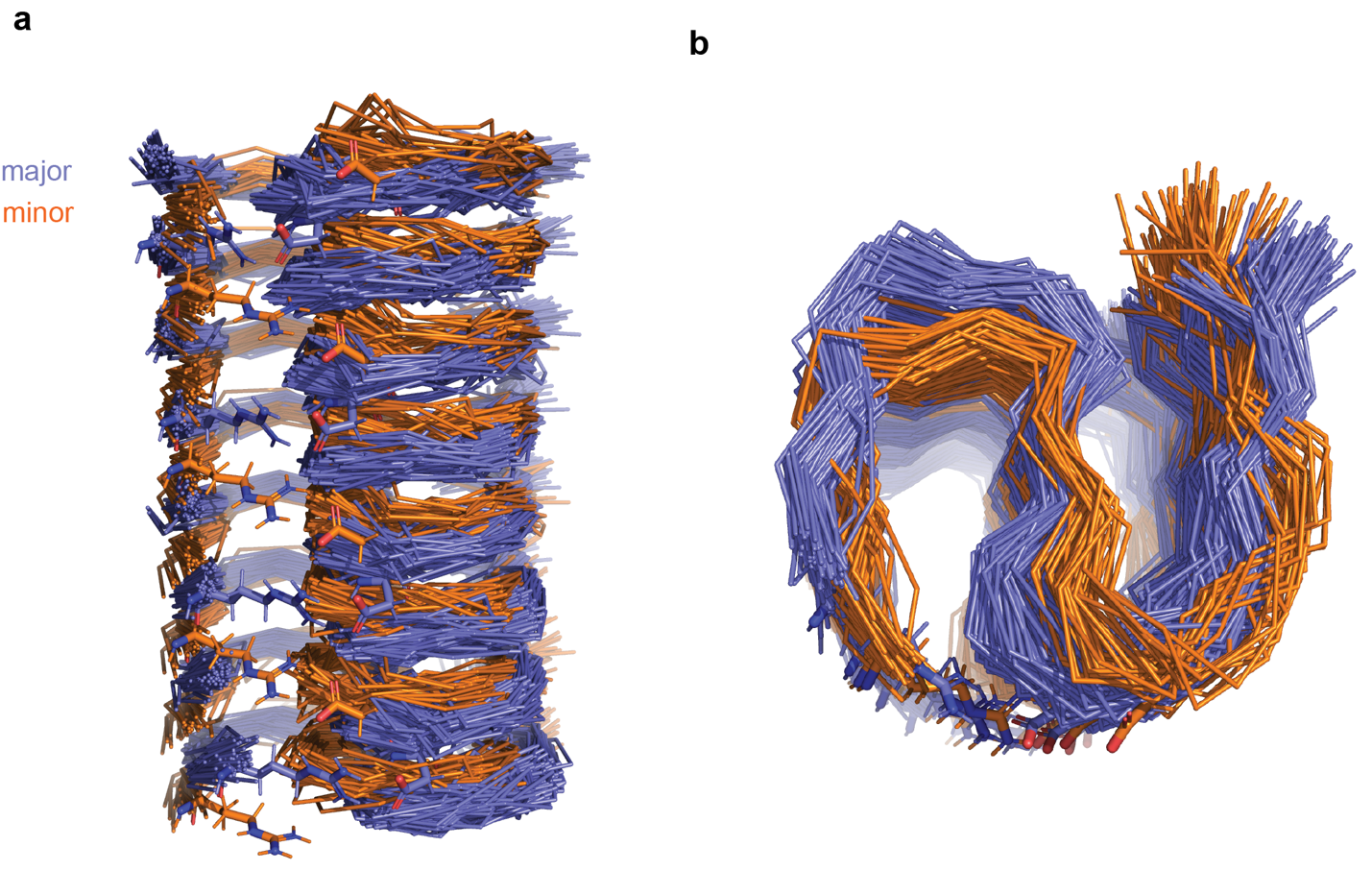

**Figure S9.** Initial Folding Structure Ensembles. Ribbon diagram displaying two structural ensembles with the R53(M45)-D462(RIPK3) contact highlighted. The slate blue represents the major species, while the orange represents the minor species.

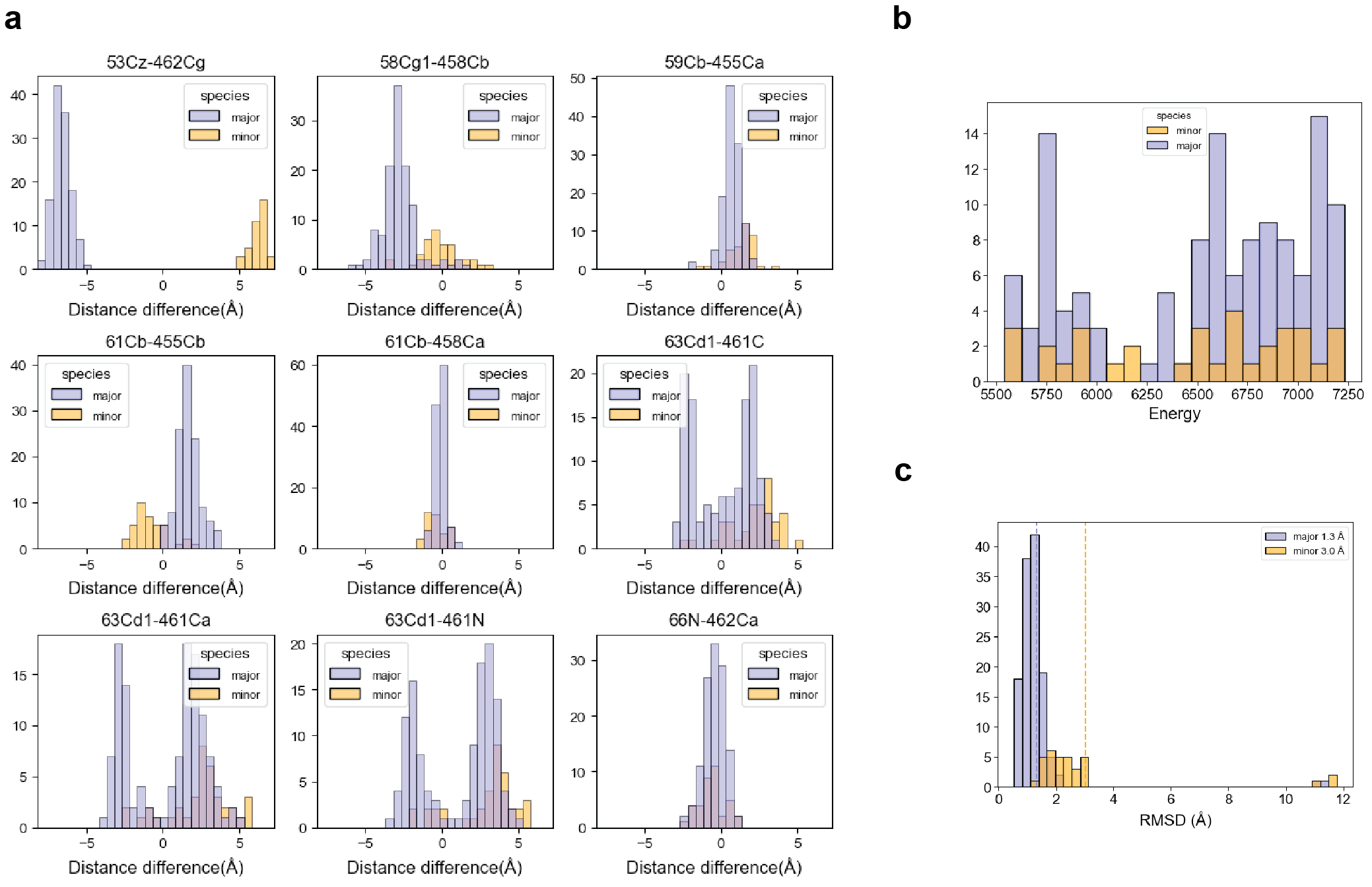

**Figure S10**. Statistics of initial folding when M45 is unambiguous in intermolecular contacts. (a) The histograms depicting the variations in distances between BA and BC contacts for each unambiguous M45-RIPK3 cross peak (as referred to in Fig. 3(a)) within the 160 lowest energy structures selected from the initial set of 320 calculated structures in the initial folding. A distribution centered around zero signifies no preference for either BA or BC contacts, while a left-shifted distribution suggests a preference for BA contact, and a right-shifted distribution indicates a preference for BC contact. The slate blue represents the major species, while the orange represents the minor species. (b) The energy distribution of the 160 structures. (c) The distribution of the RMSDs of alignment to the lowest energy structure.

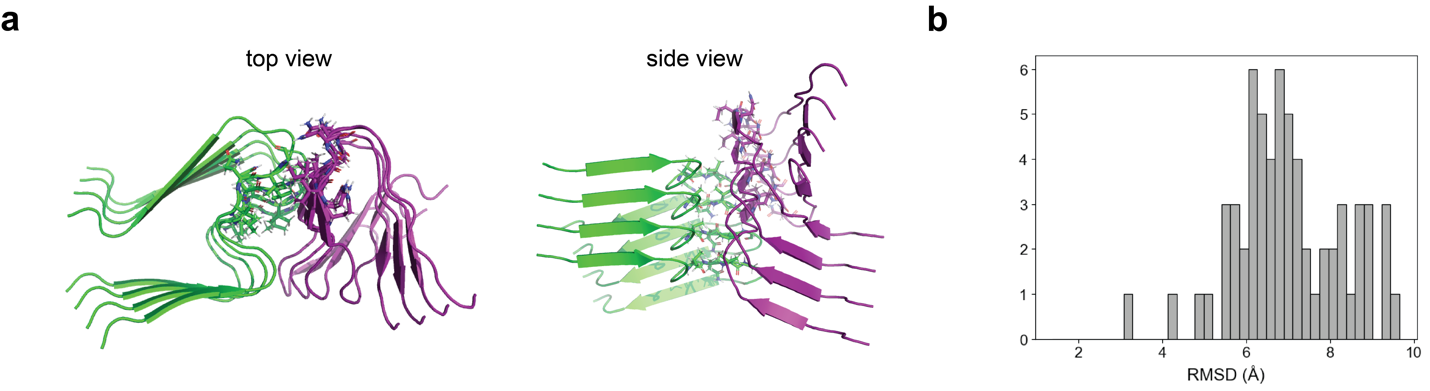

**Figure S11**. Initial folding results for scheme III. (a) The best structure among 320 calculated structures. (b) RMSM distribution of backbone heavy atom alignments for the 160 lowest-energy structures relative to the best structure.

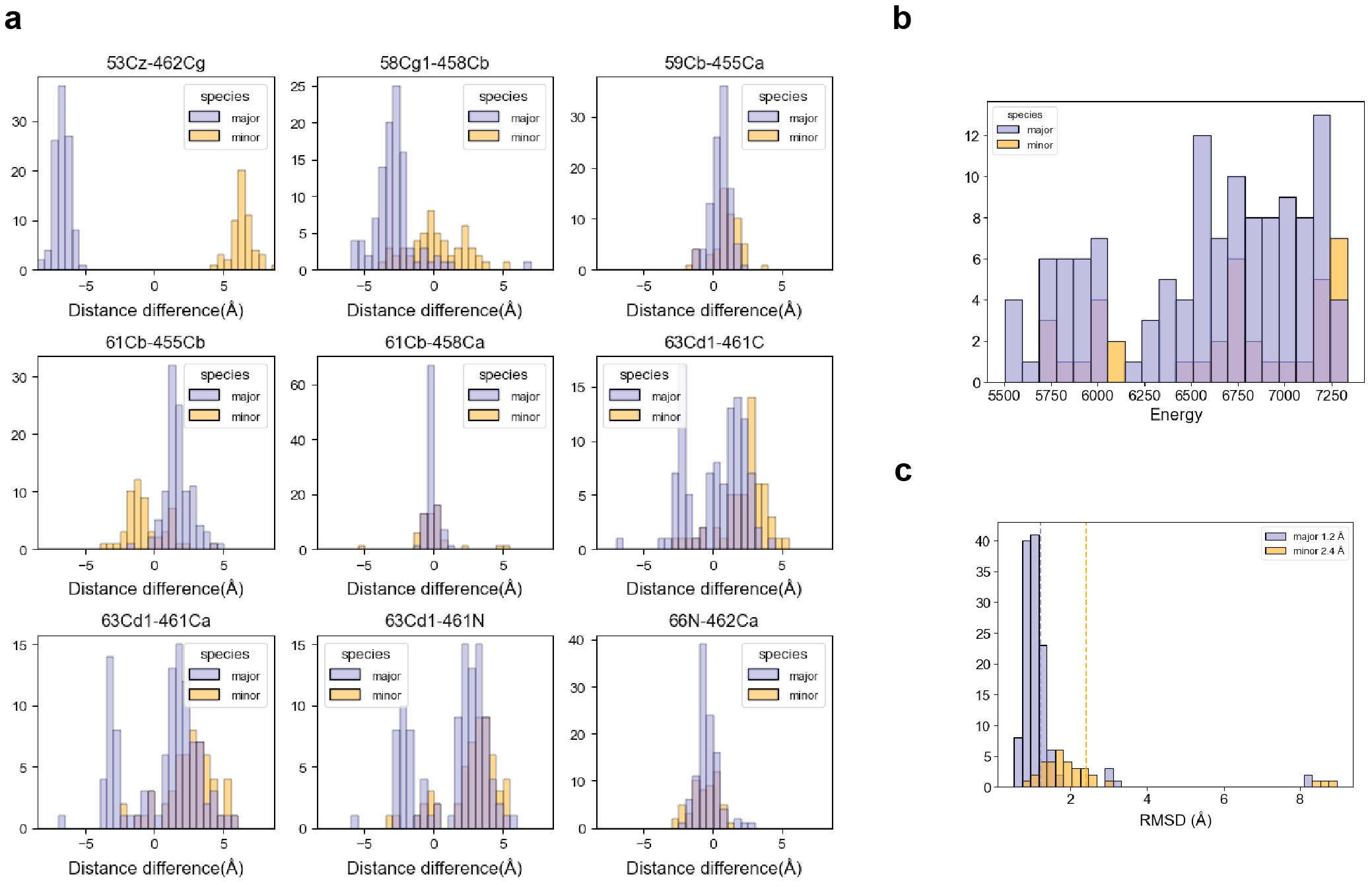

**Figure S12.** Statistics of initial folding when RIPK3 is unambiguous in intermolecular contacts. Same as **Fig. S10**.

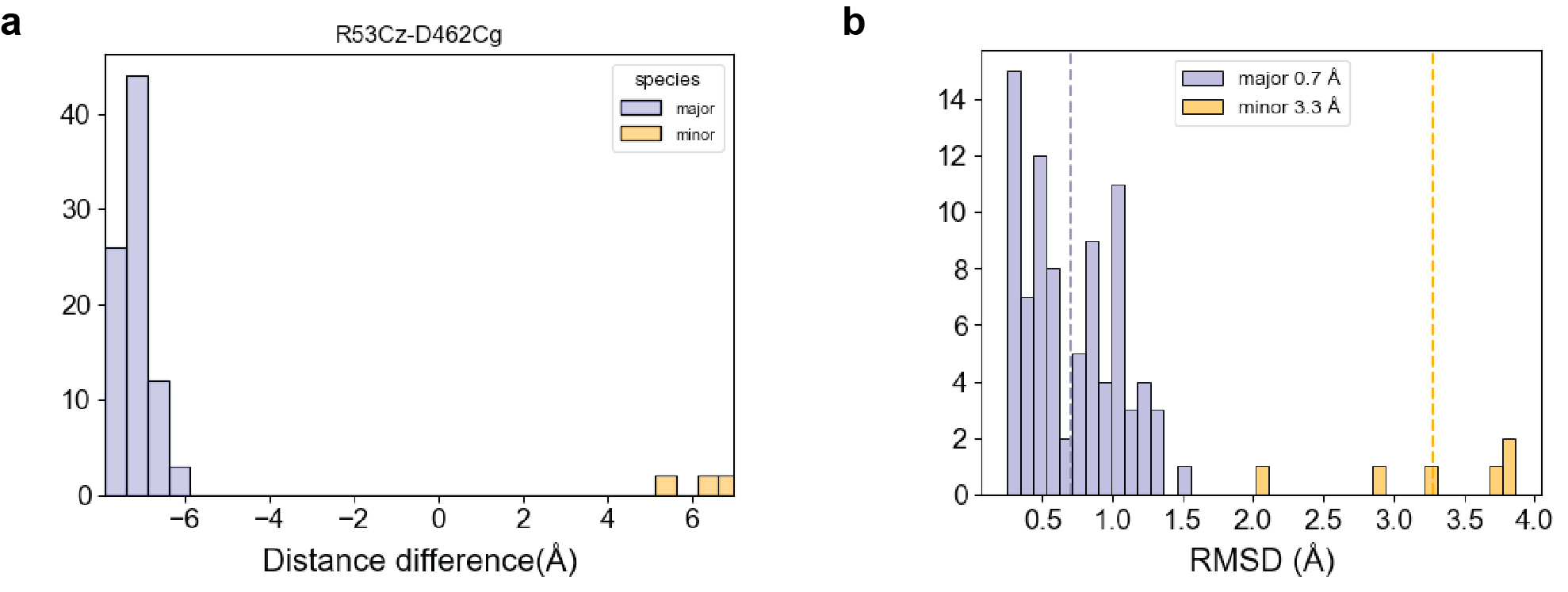

**Figure S13.** Refinement Statistics. (A) Distribution of major and minor structures among the top 90 refined models. The definitions of major and minor structures are consistent with those described previously. (B) Distribution of RMSD values for the top 90 structures when aligned to the lowest-energy structure.

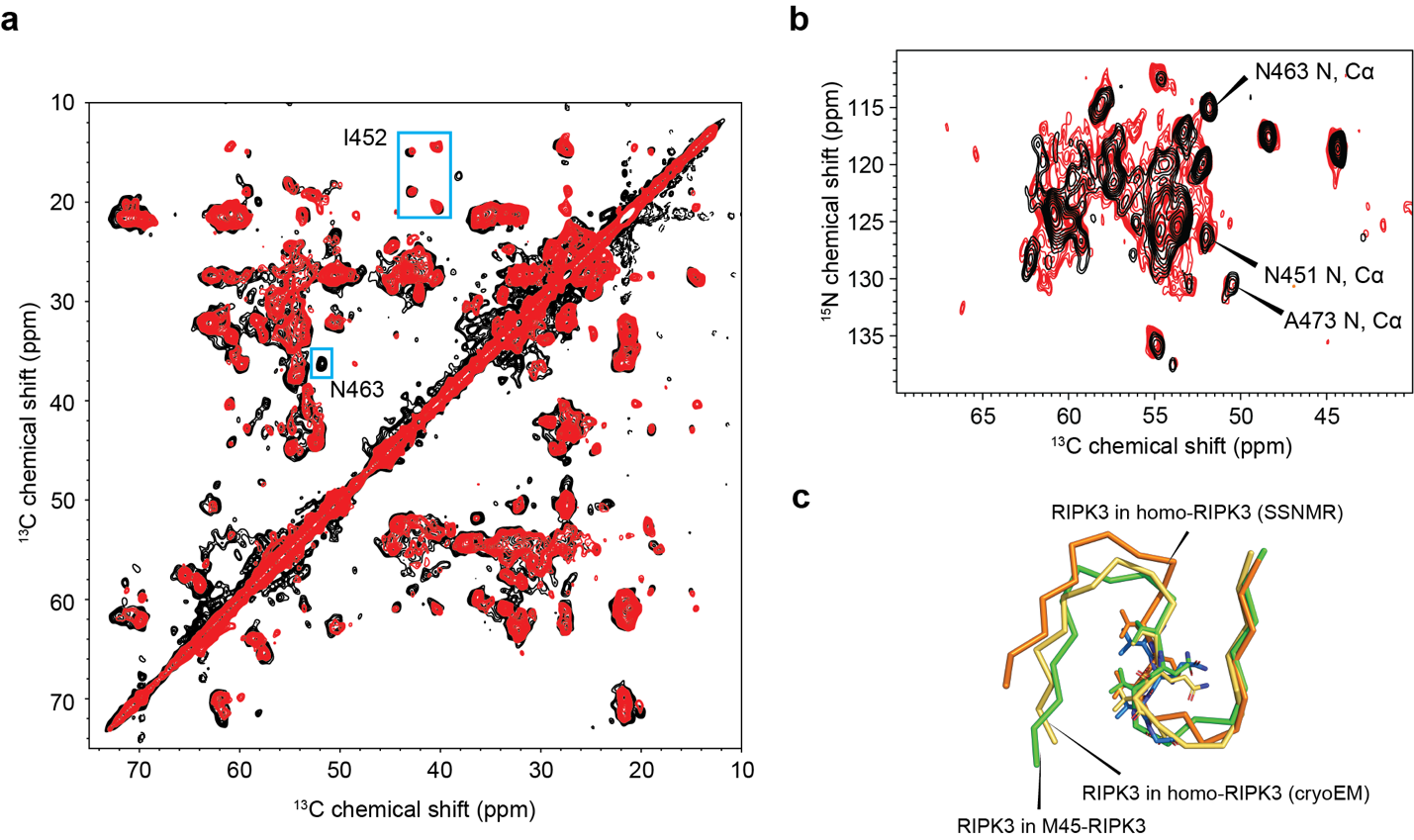

**Figure S14.** Comparison between RIPK3(homo) and RIPK3(M45). (a) and (b) show the DARR and NCA spectra comparisons between RIPK3(homo) in black and RIPK3(M45) in red, respectively. (c) is the structural alignment: green represents RIPK3(M45), yellow represents RIPK3(homo, cryo-EM) (7da4), and orange represents RIPK3(homo, SSNMR) (7dac). Backbone heavy atoms are aligned. Residues 457-469 of RIPK3(homo, SSNMR) (7dac) are aligned to the same region in RIPK3(M45), with an RMSD of 1.5 Å. Residues 450-469 of RIPK3(homo, cryo-EM) are aligned to the same region in RIPK3(M45), with an RMSD of 1.8 Å.

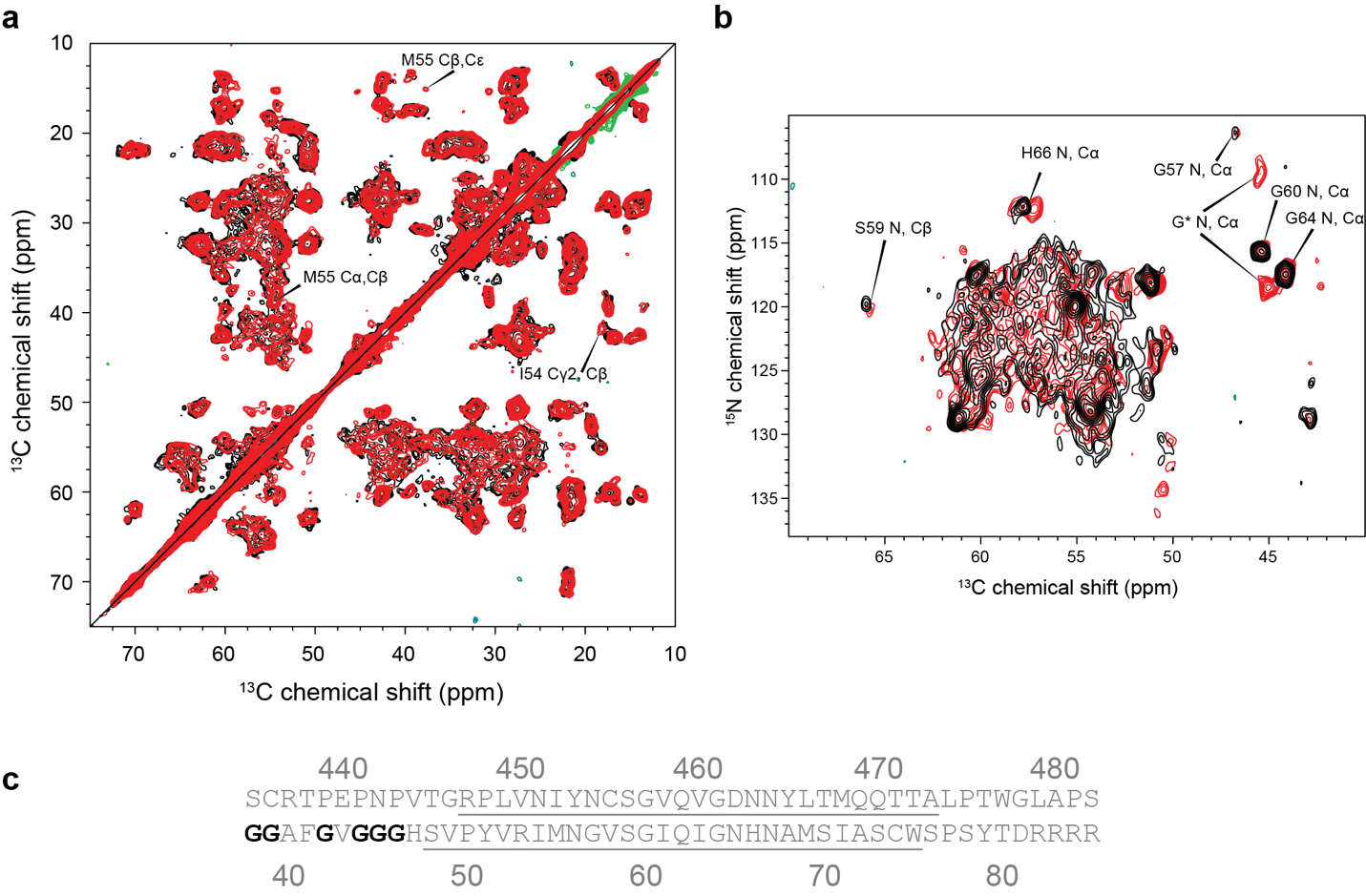

**Figure S15.** Comparison between M45(homo) in black and M45(RIPK3) in red. (a) and (b) show the DARR and NCA spectra comparisons between M45(homo) and M45(RIPK3), respectively. (c) Sequence alignment of M45 and RIPK3. Assigned residues are underlined. Several Gly residues before the N-terminal of the assigned residues in M45 are highlighted in bold black.

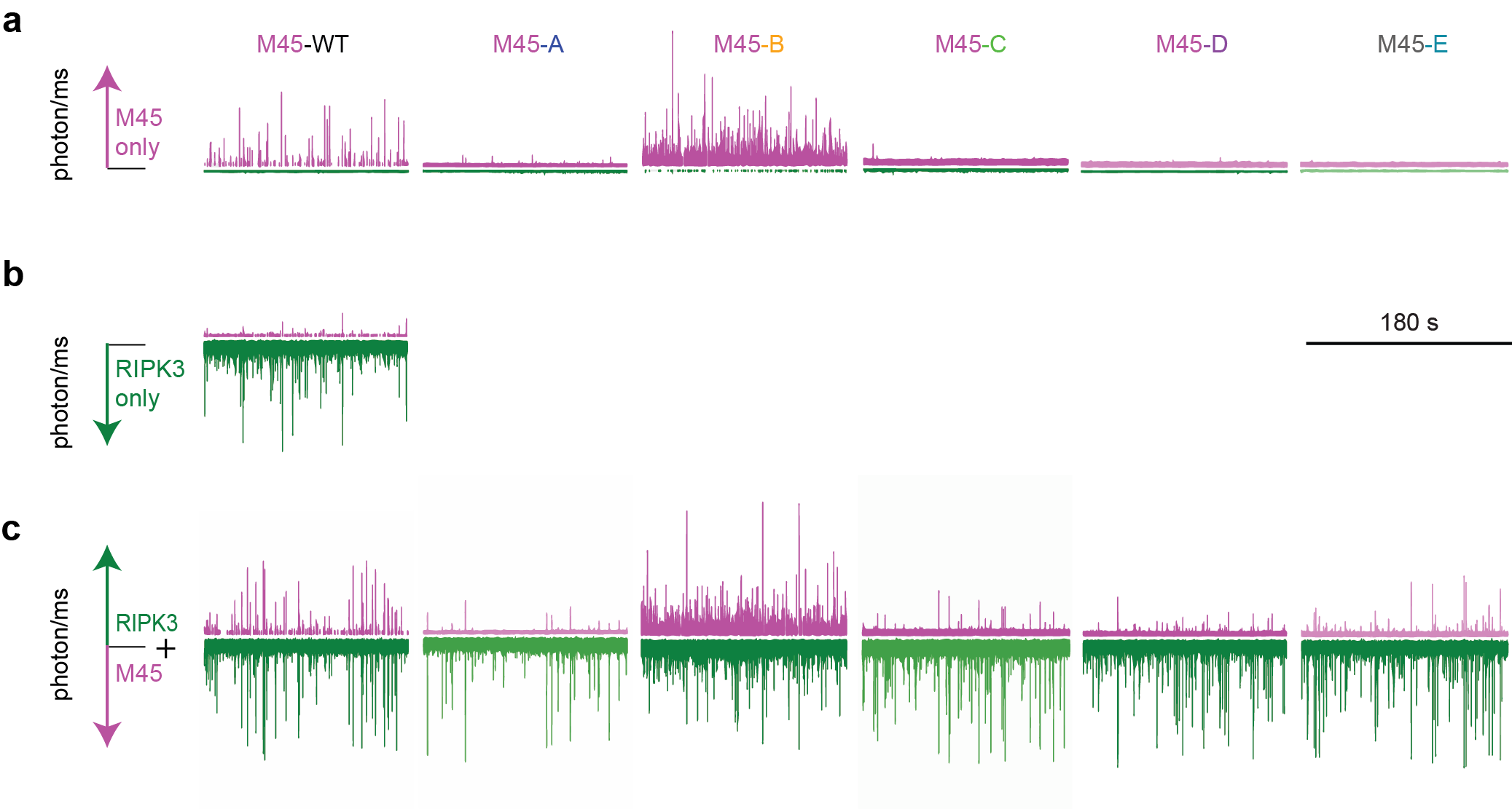

**Figure S16.** M45 homomeric assembly and heteromeric assembly with RIPK3. Full 180s traces. Simultaneous detection of mCherry (positive y-axis) and YPet (negative y-axis) fluorescence from homomeric samples of (a) WT or AAAA mutants of mCherry-M45 or (b) YPet-RIPK3 (middle panel). Samples contained only one type of protein but a small amount of bleed through YPet fluorescence was detected in the mCherry channel when very large YPet-RIPK3-containing assemblies passed through the confocal volume. (c) Simultaneous detection of fluorescence from mCherry-tagged WT and mutant M45 and YPet-RIPK3 in samples containing both RHIM proteins.

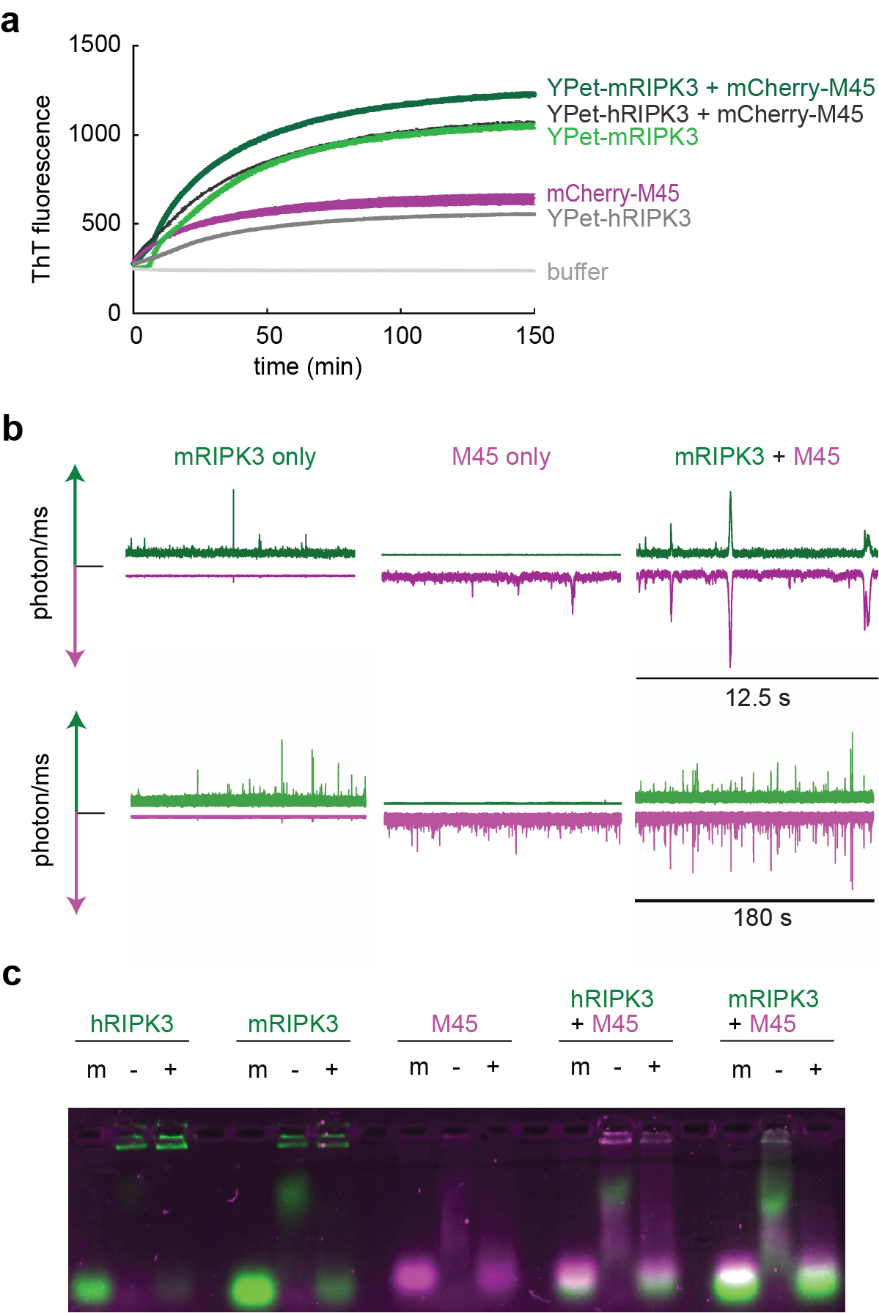

**Figure S17.** Amyloid assembly by mRIPK3 and heteromeric assembly with M45. (a) Thioflavin T fluorescence assay of amyloid assembly by YPet-mRIPK3 and hRIPK3, in the absence or presence of mCherry-M45. Fluorescence emission (arbitrary units) at 485 nm; average of triplicate values, error bars represent SD. (b) Single molecule spectroscopy with detection of YPet (positive y-axis) and mCherry (negative y-axis) fluorescence from YPet-mRIPK3 alone (left), mCherry-M45 alone (middle), or a mixture of YPet-mRIPK3 and M45 (right). Expanded representative 12.5 s sequence (upper panel) from full 180 s traces (lower panel). In YPet-RIPK3 homomeric sample, peaks detected in the mCherry channel arise from bleed-through. (c) SDS semi-denaturing agarose gel electrophoresis indicates the nature of species observed when hRIPK3 or mRIPK3, and WT M45, are maintained individually in urea (m), or incubated alone or together under assembly-permissive conditions (-) and then treated with 2% SDS (+).

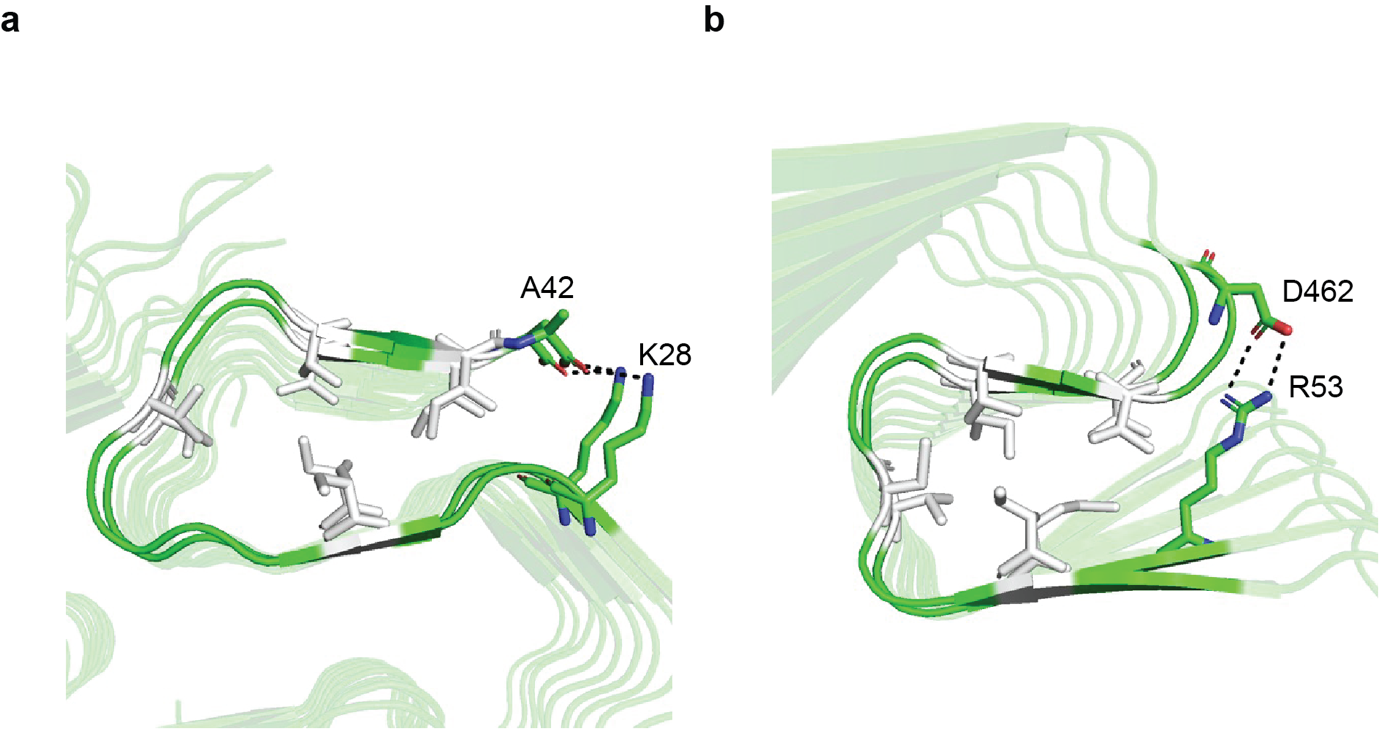

**Figure S18**. Comparison between the hydrophobic cores sealed by a salt bridge. Hydrophobic bulky residues are colored in white and hydrogen bonds are indicated by black dotted lines. (a) The hydrophobic core of Aβ42(5KK3) C-terminal tail, tethered by K28 and the terminal carboxylate of A42. (b) The hydrophobic core of M45-RIPK3 between the first and second strand, anchored by R53 and D462.

**Table S1.** Probabilities of β-sheet

| M45  residue | M45  probability | M45 cross peak | RIPK3  residue | RIPK3  probability | RIPK3 cross peak |
| --- | --- | --- | --- | --- | --- |
| S48 | N/A | * | R447 | 22.8 | Ca-Cb |
| V49 | ** | Ca-Cb | P448 | 58.4 | Ca-Cb |
| P50 | 58.7 | Ca-Cb | L449 | N/A | * |
| Y51 | 76.7 | Ca-Cb | V450 | 78.0 | Ca-Cb |
| V52 | 73.5 | Ca-Cb | N451 | 91.1 | Ca-Cb |
| R53 | 87.5 | Ca-Cb | I452 | 75.4 | Ca-Cb |
| I54 | 89.5 | Ca-Cb | Y453 | 87.5 | Ca-Cb |
| M55 | 91.3 | Ca-Cb | N454 | 72.2 | Ca-Cb |
| N56 | 60.2 | Ca-Cb | C455 | 11.5 | Ca-Cb |
| G57 | 91.4 | Ca-C | S456 | 73.6 | Ca-Cb |
| V58 | 86.4 | Ca-Cb | G457 | ** | Ca-C |
| S59 | 78.5 | Ca-Cb | V458 | 82.7 | Ca-Cb |
| G60 | 4.5 | Ca-C | Q459 | 93.8 | Ca-Cb |
| I61 | 92.1 | Ca-Cb | V460 | 62.2 | Ca-Cb |
| Q62 | 86.5 | Ca-Cb | G461 | 81.7 | Ca-C |
| I63 | 87.9 | Ca-Cb | D462 | 89.3 | Ca-Cb |
| G64 | 81.4 | Ca-Cb | N463 | 19.1 | Ca-Cb |
| N65 | 87.5 | Ca-Cb | N464 | 38.0 | Ca-Cb |
| H66 | 15.7 | Ca-C | Y465 | 88.1 | Ca-C |
| N67 | 45.6 | Ca-Cb | L466 | 67.5 | Ca-Cb |
| A68 | 84.4 | Ca-Cb | T467 | 54.4 | Ca-Cb |
| M69 | 89.5 | Ca-Cb | M468 | 51.2 | Ca-Cb |
| S70 | 52.1 | Ca-Cb | Q469 | 71.4 | Ca-Cb |
| I71 | 74.5 | Ca-Cb | Q470 | 79.7 | Ca-Cb |
| A72 | 61.6 | Ca-Cb | T471 | 74.1 | Ca-Cb |
| S73 | 56.8 | Ca-Cb | T472 | 65.8 | Ca-Cb |
| C74 | N/A | * | A473 | 93.5 | Ca-Cb |
| W75 | 41.2 | Ca-Cb |  |  |  |

Notes: Ca-Cb or Ca-C cross peaks are used for inference. * indicates insufficient data due to incomplete assignment, while ** denotes unconventional chemical shifts and failure to infer from the database.

**Table S2.** Unambiguous intermolecular and intramolecular distance constraints

| M45 intramolecular | | RIPK3 intramolecular | | M45-RIPK3 intermolecular | |
| --- | --- | --- | --- | --- | --- |
| Long-range  (i-j>4) | Medium-range  (i-j≤4) | Long-range  (i-j>4) | Medium-range (i-j≤4) | d > 4 | d ≤4 |
| S59C-M69Ce | P50Cd-V52Ca | I452Cd1-G457C | R447Ca-L449Cd1 | R53Cz-D462Cg | V58Cg2-V458Cb |
| S59Cb-M69Ce | R53Cb-M55Cb | I452Cg1-G457C | C455Ca-G457C | R53Cd-V460Ca * | S59Cb-C455Ca |
| S59Cb-I71Cg1 | I54Cg1-N56Cb | I452Cb-V458Cb | C455Cb-V458Ca | Q62Cb-L466Cb * | I61Cb-C455Cb |
| S59Cb-I71Cd1 | G57C-S59Cb | S456C-M468Ce | G457C-Q459Ca |  | I61Cb-V458Ca |
| G60Ca-M69Ce | V58Cg*-I61Cb | S456Cb-M468Ce | G457C-459Cg |  | I63Cd1-G461Ca |
| G60N-I71Cd1 | S59Ca-I61Cg2 | S457Ca-L466Cb | V458Cb-460Ca |  | I63Cd1-G461C |
| Q62Cd-M69Ce | S59Cb-I61Cg2 | G457Ca-L466Cg | Y465C-T467Cb |  | A68Cb-T467Cb * |
| Q62Cg-M69Ce | I61Cd1-I63Cg1 | G457C-M468Ce | L466Cb-M468Cg |  | I71Ca-Q469Cb * |
|  | Q62Ca-G64C | Q459Cd-L466Cb | L466Cd1-M468Ce |  | N65Ca-D462Ca * |
|  | Q62Cb-G64C | Q459Cd-L466Cg | T467Cb-Q469Cd |  | A72Ca-M468Ca * |
|  | A68Cb-S70Cb |  | Q470C-T472Cb |  | I63Cd1-G461N |
|  | M69Cb-I71Ca |  |  |  | H66N-D462Ca |
|  | I71Cg1-S73Cb |  |  |  |  |

Notes: Contacts derived from NC heteronuclear experiments (TEDOR) are underlined, while those derived from CC homonuclear experiments (DARR) are not underlined. * indicates contacts obtained by comparing experimental spectra with back-predicted contact spectra of the initial structure and they are only used in the refinement .

**Table S3.** Low-ambiguity intermolecular and intramolecular distance constraints

| M45 intramolecular | RIPK3 intramolecular | M45-RIPK3 intermolecular |
| --- | --- | --- |
| I54Ca/V58Ca-N56Cb | I452Cb/N464Cb-V458Cb | 62GlnCg/460ValCb-69MetCe |
| Q62Ca/M55Ca-I61Cg2 | G457C/G461C-Q459Ca | 62GlnCg/460ValCb-61IleCg2 |
| M55Ce/M69Ce-S59Cb | G457C/G461C-Q459Cb | 460ValCa/452IleCa-55MetCe |
| I61Cd1/M55Ce-V58Cg1 | G457C/G461C-Q459Cg | 458ValCg1/460ValCg1-58ValCb |
|  | P448Cg/I452Cg1-C455Ca | 62GlnCa/55MetCa-466LeuCda |
|  |  | 62GlnCg/460ValCb-61IleCg1 |
|  |  | 69MetCg/55MetCg-452IleCg2 |

**Table S4.** Sequential contacts

| M45 | RIPK3 |
| --- | --- |
| S48CB-V49CB | R447CD-P448CD |
| S48CB-V49CG1 | P448CA-L449CD1 |
| S48CB-V49CG2 | P448CA-L449CD2 |
| Y51C-V52N | P448CD-L449CD1 |
| Y51CA-V52N | P448CD-L449CD2 |
| V52C-R53N | N451C-I452N |
| V52CA-R53N | N451C-I452CB |
| R53CA-I54CD1 | N451CA-I452N |
| I54CA-M55CA | N451CA-I452CG1 |
| I54CA-M55CG | N451CA-I452CG2 |
| I54CA-M55CE | N451CB-I452N |
| M55CG-N56CB | N451CB-I452CA |
| N56CA-G57C | N451CB-I452CB |
| G57C-V58N | I452C-Y453CA |
| G57C-V58CA | I452CB-Y453CA |
| G57C-V58CB | N454CG-C455CA |
| G57C-V58CG1 | C455C-S456CA |
| V58C-S59N | C455CA-S456N |
| V58CA-S59N | C455CB-S456N |
| V58CB-S59N | C455CB-S455CB |
| V58CG1-S59N | S456C-G457CA |
| V58CG1-S59CB | S456CA-G457N |
| V58CG2-S59CB | S456CA-G457CA |
| S59C-G59N | S456CB-G457C |
| S59C-G59CA | S456CB-G457CA |
| S59CA-G59N | G457C-V458N |
| S59CB-G59N | G457C-V458CA |
| S59CB-G59CA | G457C-V458CB |
| G60C-I61N | G457CA-V458C |
| G60CA-I61N | G457CA-V458CA |
| G60CA-I61CA | G457CA-V458CB |
| G60CA-I61CB | V458C-Q459N |
| G60CA-I61CG1 | V458N-Q459CD |
| G60CA-I61CG2 | V458CA-Q459N |
| I61C-Q62N | V458CB-Q459N |
| I61CA-Q62CA | Q459C-V460N |
| I61CB-Q62CB | V460C-G461CA |
| I61CG1-Q62CA | V460CA-G461CA |
| I61CG1-Q62CG | V460CB-G461CA |
| I61CG2-Q62CA | V460CA-G461C |
| I61CG2-Q62CB | V460CB-G461C |
| I61CG2-Q62CG | V460CA-G461N |
| Q62CA-I63CA | V460CB-G461N |
| Q62CG-I63CA | G461C-D462CA |
| Q62CG-I63CG2 | G461C-D462CB |
| I63CA-G64N | D462C-N463N |
| I63CB-G64CA | D462C-N463CA |
| I63CG1-G64C | D462CA-N463CB |
| G64C-N65N | D462CB-N463CB |
| G64C-N65CA | N463C-N464N |
| G64C-N65CB | N463C-N464CA |
| G64CA-N65N | N463CA-N464CB |
| G64CA-N65CA | N463CB-N464CA |
| N65C-H66N | Y465CA-L466CG |
| N65CA-H66N | L466CA-T467N |
| N67C-A68N | L466CB-T467N |
| N67CA-A68N | L466CG-T467N |
| A68C-M69N | L466CA-T467C |
| A68CA-M69N | L466CB-T467C |
| A68CA-M69CB | L466CG-T467C |
| A68CB-M69N | L466CG-T467CA |
| M69CB-S70CB | T467C-M468N |
| S70C-I71N | T467C-M468CA |
| S70CA-I71N | T467CA-M468N |
| I71C-A72N | M468CA-Q469N |
| I71CA-A72N | M468CB-Q469N |
| I71CG2-A72N | M468CB-Q469CD |
| A72CB-S73CB | Q470CA-T471N |
| S73CB-C74CB | Q470CB-T471N |
|  | Q470CG-T471N |
|  | Q470CB-T471CA |
|  | T471CA-T472N |
|  | T471CB-T472N |
|  | T472CA-A473N |
|  | T472CB-A473N |

**Table S5.** Backbone dihedral angle and inter-strand distance constraints

| M45  resid | $\Phi$ | $\Delta\Phi$ | $\Psi$ | $\Delta\Psi$ | RIPK3 resid | $\Phi$ | $\Delta\Phi$ | $\Psi$ | $\Delta\Psi$ | Inter-strand  C-C NOE |
| --- | --- | --- | --- | --- | --- | --- | --- | --- | --- | --- |
| S48 |  |  |  |  |  |  |  |  |  |  |
| V49 |  |  |  |  |  |  |  |  |  |  |
| P50 |  |  |  |  | R447 |  |  |  |  |  |
| Y51 | -125.2 | 54.3 | 149.7 | 30.3 | P448 |  |  |  |  | 4.75 ± 0.1 |
| V52 | -126.4 | 31.2 | 149.3 | 37.1 | L449 |  |  |  |  | 4.75 ± 0.1 |
| R53 | -129.8 | 48.2 | 135.4 | 33.6 | V450 | -117.0 | 46.0 | 139.1 | 25.2 | 4.75 ± 0.1 |
| I54 | -120.7 | 36.7 | 131.0 | 41.9 | N451 | -123.1 | 45.9 | 140.4 | 25.9 | 4.75 ± 0.1 |
| M55 | -136.4 | 46.6 | 150.5 | 38.1 | I452 | -113.6 | 46.2 | 129.7 | 33.5 | 4.75 ± 0.1 |
| N56 |  |  |  |  | Y453 |  |  |  |  | 4.75 ± 0.1 |
| G57 |  |  |  |  | N454 |  |  |  |  | 4.75 ± 0.1 |
| V58 | -122.0 | 50.3 | 154.7 | 39.9 | C455 | -138.5 | 57.5 | 156.2 | 32.3 |  |
| S59 | -132.1 | 48.5 | 146.6 | 39.2 | S456 | -122.9 | 76.4 | 137.2 | 51.1 | 4.75 ± 0.1 |
| G60 |  |  |  |  | G457 |  |  |  |  |  |
| I61 | -129.5 | 40.4 | 134.9 | 28.2 | V458 | -128.5 | 32.7 | 143.1 | 42.1 | 4.75 ± 0.1 |
| Q62 | -127.1 | 42.3 | 134.4 | 42.9 | Q459 | -125.2 | 45.3 | 136.9 | 34.4 | 4.75 ± 0.1 |
| I63 | -125.0 | 32.2 | 140.8 | 42.9 | V460 | -108.0 | 40.5 | 130.4 | 27.8 | 4.75 ± 0.1 |
| G64 |  |  |  |  | G461 |  |  |  |  | 4.75 ± 0.1 |
| N65 | -122.2 | 72.2 | 168.6 | 39.2 | D462 |  |  |  |  | 4.75 ± 0.1 |
| H66 |  |  |  |  | N463 |  |  |  |  |  |
| N67 | -102.3 | 41.6 | -5.1 | 45.6 | N464 |  |  |  |  |  |
| A68 | -125.3 | 57.6 | 139.7 | 40.6 | Y465 | -109.1 | 60.6 | 121.7 | 35.2 | 4.75 ± 0.1 |
| M69 | -136.7 | 34.3 | 138.4 | 34.7 | L466 | -109.4 | 41.2 | 127.1 | 24.4 | 4.75 ± 0.1 |
| S70 | -114.6 | 39.8 | 135.8 | 41.5 | T467 | -124.9 | 49.5 | 135.5 | 50.9 | 4.75 ± 0.1 |
| I71 |  |  |  |  | M468 | -110.6 | 48.8 | 128.4 | 46.3 | 4.75 ± 0.1 |
| A72 | -137.5 | 50.2 | 153.4 | 35.5 | Q469 | -100.2 | 39.7 | 134.6 | 30.7 | 4.75 ± 0.1 |
| S73 | -118.5 | 56.2 | 141.4 | 59.5 | Q470 | -124.8 | 35.5 | 147.4 | 44.3 | 4.75 ± 0.1 |
| C74 |  |  |  |  | T471 |  |  |  |  |  |
| W75 |  |  |  |  | T472 |  |  |  |  |  |
|  |  |  |  |  | A473 |  |  |  |  |  |

**Table S6.** Hydrogen bond constraints

| Donor | Acceptor |
| --- | --- |
| R53HN1 | D462OD1 |
| R53HN2 | D462OD2 |

**Table S7.** Chemical shift assignment of M45

| M45 resid | C | Ca | Cb | Cd*/Cd/Cd1 | Ce/Ce1/Ce2 | Ce3 | Cg/Cg1 | Cg2 | Ch2/Cz | N | Nd2/Ne2 |
| --- | --- | --- | --- | --- | --- | --- | --- | --- | --- | --- | --- |
| Ser48 |  |  | 65.5 |  |  |  |  |  |  |  |  |
| Val49 |  | 59.3 | 37.7 |  |  |  |  |  |  |  |  |
| Pro50 | 172.9 | 62.7 | 32.3 | 50.5 |  |  | 27.2 |  |  | 130.8 |  |
| Tyr51 | 175.3 | 56.0 | 40.3 |  |  |  | 129.5 |  |  |  |  |
| Val52 |  | 61.0 | 35.3 |  |  |  |  |  |  | 115.2 |  |
| Arg53 |  | 56.0 | 34.9 | 43.3 |  |  | 27.9 |  | 158.5 | 123.1 |  |
| Ile54 | 174.6 | 60.5 | 41.9 | 13.6 |  |  | 28.5 | 18.2 |  | 126.6 |  |
| Met55 | 174.3 | 54.2 | 37.6 |  | 15.3 |  | 30.7 |  |  |  |  |
| Asn56 | 175.0 | 53.6 | 41.3 |  |  |  | 177.4 |  |  | 119.9 |  |
| Gly57 | 170.8 | 46.5 |  |  |  |  |  |  |  | 104.8 |  |
| Val58 | 174.7 | 60.3 | 35.7 |  |  |  | 22.1 | 21.6 |  | 116.3 |  |
| Ser59 | 172.7 | 55.2 | 65.8 |  |  |  |  |  |  | 118.6 |  |
| Gly60 | 174.3 | 45.3 |  |  |  |  |  |  |  | 114.4 |  |
| Ile61 | 173.9 | 61.0 | 42.9 | 14.9 |  |  | 28.9 | 16.6 |  | 127.4 |  |
| Gln62 | 174.0 | 54.2 | 32.1 | 177.8 |  |  | 33.4 |  |  | 127.0 | 113.9 |
| Ile63 | 174.3 | 60.0 | 42.6 | 13.8 |  |  | 27.4 | 17.2 |  | 124.3 |  |
| Gly64 | 171.1 | 44.0 |  |  |  |  |  |  |  | 116.0 |  |
| Asn65 | 175.1 | 51.3 | 42.5 |  |  |  | 176.9 |  |  | 116.9 | 114.7 |
| His66 | 175.5 | 57.7 |  | 123.0 | 137.9 |  |  |  |  | 111.2 |  |
| Asn67 | 175.0 | 52.8 | 39.9 |  |  |  | 175.9 |  |  |  |  |
| Ala68 |  | 50.6 | 22.1 |  |  |  |  |  |  | 122.6 |  |
| Met69 |  | 54.4 | 38.6 |  | 15.3 |  | 30.7 |  |  | 126.3 |  |
| Ser70 |  | 55.1 | 64.4 |  |  |  |  |  |  |  |  |
| Ile71 | 175.9 | 60.2 | 39.9 | 13.9 |  |  | 27.7 | 17.4 |  | 125.7 |  |
| Ala72 |  | 51.0 | 21.1 |  |  |  |  |  |  | 122.8 |  |
| Ser73 | 173.1 | 55.1 | 64.7 |  |  |  |  |  |  |  |  |
| Trp75 | 174.9 | 57.8 | 30.3 |  | 138.2 | 119.8 | 111.2 |  | 123.7 |  |  |

**Table S8.** Chemical shift assignment of RIPK3

| RIPK3 resid | C | Ca | Cb | Cd1/Cd*/Cd | Ce/Ce* | Cg1/Cg | Cg2 | Cz | N | Ne2 |
| --- | --- | --- | --- | --- | --- | --- | --- | --- | --- | --- |
| Arg447 |  | 56.7 | 30.3 | 43.5 |  |  |  | 159.5 |  |  |
| Pro448 |  | 62.7 | 32.2 | 50.3 |  | 27.4 |  |  | 136.4 |  |
| Leu449 |  |  |  | 24.9 |  |  |  |  |  |  |
| Val450 |  | 60.3 | 35.1 |  |  |  |  |  |  |  |
| Asn451 | 173.5 | 52.0 | 42.9 |  |  |  |  |  | 125.1 |  |
| Ile452 | 173.7 | 61.0 | 40.3 | 14.5 |  | 27.6 | 20.7 |  | 122.6 |  |
| Tyr453 | 122.7 | 56.3 | 40.5 | 133.9 | 117.4 |  |  | 157.1 |  |  |
| Asn454 |  | 53.3 | 41.0 |  |  | 179.9 |  |  |  |  |
| Cys455 | 174.2 | 58.4 | 32.0 |  |  |  |  |  | 114.1 |  |
| Ser456 | 173.0 | 57.6 | 65.5 |  |  |  |  |  | 118.2 |  |
| Gly457 | 170.3 | 48.4 |  |  |  |  |  |  | 116.4 |  |
| Val458 | 174.2 | 59.9 | 36.3 |  |  | 21.3 | 21.9 |  | 123.8 |  |
| Gln459 | 174.5 | 53.7 | 32.3 | 177.5 |  | 33.8 |  |  | 124.3 | 112.9 |
| Val460 | 174.5 | 60.9 | 33.5 |  |  | 21.3 |  |  | 123.3 |  |
| Gly461 | 170.4 | 44.5 |  |  |  |  |  |  | 117.4 |  |
| Asp462 | 175.5 | 52.4 | 44.4 |  |  | 181.1 |  |  | 119.6 |  |
| Asn463 | 173.9 | 51.9 | 36.4 |  |  |  |  |  | 113.7 |  |
| Asn464 | 173.9 | 53.4 | 40.2 |  |  | 177.0 |  |  | 115.8 |  |
| Tyr465 | 173.4 | 56.6 |  |  | 117.7 |  |  |  |  |  |
| Leu466 | 174.6 | 55.0 | 44.8 | 24.5 |  | 28.7 |  |  | 134.6 |  |
| Thr467 | 172.3 | 57.9 | 70.8 |  |  |  | 20.0 |  | 120.4 |  |
| Met468 | 174.6 | 54.6 | 34.3 |  | 16.6 | 32.2 |  |  | 124.5 |  |
| Gln469 | 175.0 | 54.7 | 30.7 | 179.4 |  |  |  |  | 123.9 |  |
| Gln470 | 175.2 | 53.0 | 33.4 |  |  | 37.6 |  |  |  |  |
| Thr471 |  | 61.6 | 71.2 |  |  |  |  |  | 117.6 |  |
| Thr472 |  | 62.2 | 70.4 |  |  |  |  |  | 126.5 |  |
| Ala473 |  | 50.5 | 23.5 |  |  |  |  |  | 129.2 |  |

**Table S9.** Table of 2D experiments

| Dataset  _ID | Sample | Experiment | Acq_ td_f1 | Acq_ td_f2 | Acq_ time_ f1 | Acq_ time_ f2 | Proc_f1 | Proc_f2 | Proc_ td_f1 | Proc_ td_f2 | Powers | Instrument  (B_0_ / probe) | Experiment Condition (spinning / set temp / actual temp) | S/N |
| --- | --- | --- | --- | --- | --- | --- | --- | --- | --- | --- | --- | --- | --- | --- |
| 1 | uM45_naRIPK3 | DARR 75ms | 800 | 1600 | 8.5 | 17.0 | qsine:3 | qsine:3 | 800 | 800 | ^1^H 240W 2.5us ^13^C 80W 4.4us | 900MHz /  3.2mm Efree | 16kHz /  273K / 280K | 10.2 |
| 2 | uM45_naRIPK3 | DARR 350ms | 500 | 2048 | 5.3 | 13.1 | qsine:3 | qsine:3 | 500 | 1280 | ^1^H 50W 2.45us ^13^C 80W 4.5us | 750MHz /  1.9mm HXY | 16.66 kHz /  255K / 275K | 4.9 |
| 3 | uM45_naRIPK3 | N(CO)CX 50ms | 80 | 2048 | 7.2 | 13.1 | qsine:3 | qsine:3 | 80 | 1536 | ^1^H 50W 2.25us ^13^C 80W 4.4us ^15^N 100W 4.5us | 750MHz /  1.9mm HXY | 16.66 kHz /  265K / 285K | 4.1 |
| 4 | uM45_naRIPK3 | N(CA)CX 50ms | 84 | 2560 | 7.6 | 16.4 | qsine:3 | qsine:3 | 84 | 1536 | ^1^H 50W 2.25us ^13^C 80W 4.4us ^15^N 100W 4.5us | 750MHz /  1.9mm HXY | 16.66 kHz /  265K / 285K | 5.1 |
| 5 | uM45_naRIPK3 | CA(N)CO | 120 | 2048 | 5.4 | 13.1 | gm:  -30; 0.08 | gm:  -30; 0.08 | 120 | 1536 | ^1^H 50W 2.25us ^13^C 80W 4.4us ^15^N 100W 4.5us | 750MHz /  1.9mm HXY | 33.33kHz /  250K / 285K | 2.1 |
| 6 | uM45_naRIPK3 | NCA 4.8ms | 80 | 2048 | 7.2 | 13.1 | qsine:3 | qsine:3 | LP:28 | 2048 | ^1^H 50W 2.35us ^13^C 80W 4.4us ^15^N 100W 4.5us | 750MHz /  1.9mm HXY | 16.66 kHz /  250K / 270K | 8.9 |
| 7 | naM45_uRIPK3 | DARR 75ms | 1024 | 2560 | 6.1 | 15.4 | qsine:3 | qsine:3 | 1024 | 1536 | ^1^H 240W 2.88us ^13^C 95W 3.75us | 900MHz /  3.2mm Efree | 16kHz /  273K / 280K | 12.8 |
| 8 | naM45_uRIPK3 | NCA 4.5ms | 64 | 2048 | 8.2 | 15.3 | qsine:3 | qsine:3 | 64 | 2048 | ^1^H 240W 2.75us ^13^C 90W 3.75us ^15^N 100W 4.3us | 900MHz /  3.2mm Efree | 16kHz /  273K / 280K | 9.7 |
| 9 | naM45_uRIPK3 | N(CO)CX 75ms | 64 | 2048 | 6.3 | 12.3 | qsine:3 | qsine:3 | 64 | 1280 | ^1^H 240W 2.88us ^13^C 95W 3.75us ^15^N 150W 3.8us | 900MHz /  3.2mm Efree | 16kHz /  273K / 280K | 5.6 |
| 10 | naM45_uRIPK3 | N(CA)CX 75ms | 64 | 2048 | 6.3 | 15.3 | qsine:3 | qsine:3 | 64 | 1280 | ^1^H 240W 2.88us ^13^C 95W 3.75us ^15^N 150W 3.8us | 900MHz /  3.2mm Efree | 16kHz / 273K / 280K | 5.3 |
| 11 | naM45_uRIPK3 | CA(N)CO | 116 | 2718 | 7.0 | 14.9 | qsine:3 | qsine:3 | 116 | 1024 | ^1^H 240W 2.88us ^13^C 95W 3.75us ^15^N 150W 3.8us | 900MHz /  3.2mm Efree | 16kHz / 273K / 280K | 7.4 |
| 12 | 2M45_naRIPK3 | DARR 350ms | 632 | 2048 | 5.7 | 13.1 | qsine:3 | qsine:3 | 632 | 1536 | ^1^H 50W 2.45us ^13^C 80W 4.5us | 750MHz /  1.9mm HXY | 16.66kHz / 265K / 285K | 4.2 |
| 13 | 13M45_naRIPK3 | DARR 350ms | 600 | 2048 | 6.3 | 13.1 | qsine:3 | qsine:3 | 600 | 1600 | ^1^H 50W 2.45us ^13^C 80W 4.5us | 750MHz /  1.9mm HXY | 16.66kHz / 265K / 285K | 6.5 |
| 14 | naM45_13RIPK3 | DARR 350ms | 500 | 2048 | 5.3 | 13.1 | qsine:3 | qsine:3 | 500 | 1280 | ^1^H 50W 2.4us ^13^C 80W 4.5us | 750MHz /  1.9mm HXY | 16.66kHz / 265K / 285K | 9.8 |
| 15 | naM45_2RIPK3 | DARR 350ms | 700 | 2048 | 6.3 | 13.1 | qsine:3 | qsine:3 | 700 | 1280 | ^1^H 50W 2.4us ^13^C 80W 4.5us | 750MHz /  1.9mm HXY | 16.66kHz / 265K / 285K | 8.1 |
| 16 | 13M45_2RIPK3 | DARR 500ms | 500 | 2048 | 5.3 | 13.1 | qsine:3 | qsine:3 | 450 | 1280 | ^1^H 50W 2.45us ^13^C 80W 4.5us | 750MHz /  1.9mm HXY | 16.66kHz / 250K / 270K | 8.3 |
| 17 | 2M45_3RIPK3 | DARR 500ms | 500 | 2108 | 4.5 | 13.5 | qsine:3 | qsine:3 | 500 | 1200 | ^1^H 350W 2.675us ^13^C 160W 2.8us | 900MHz /  1.6mm Efree | 20kHz / 265K / 280K | 6.4 |
| 18 | 13M45_2RIPK3 | TEDOR 7.2ms | 280 | 2186 | 8.4 | 14.0 | qsine:3 | qsine:3 | 300 | 1280 | ^1^H 50W 2.5us ^13^C 80W 4.5us ^15^N 100W 4.5us | 750MHz /  1.9mm HXY | 16.66kHz / 250K / 270K | 5.7 |
| 19 | u_homoM45 | DARR 75ms | 1200 | 1926 | 10.6 | 16.9 | qsine:3 | qsine:3 | 905 | 1024 | ^1^H 240W 2.75us ^13^C 95W 3.75us | 900MHz /  3.2mm Efree | 16kHz / 266K / 275K | 15.1 |
| 20 | u_homoM45 | NCA 6ms | 90 | 2270 | 9.9 | 16.9 | qsine:3 | qsine:3 | 90 | 1536 | ^1^H 240W 2.75us ^13^C 90W 3.75us ^15^N 100W 4.3us | 900MHz /  3.2mm Efree | 16kHz / 266K / 275K | 5.6 |
| 21 | u_homoRIPK3 | DARR 50ms | 356 | 2048 | 3.7 | 13.1 | qsine:3 | qsine:3 | LP:16 | 1536 | ^1^H 50W 2.4us ^13^C 80W 4.5us | 750MHz /  1.9mm HXY | 16.66 kHz / 255K / 275K | 13.1 |
| 22 | u_homoRIPK3 | NCA 4ms | 128 | 2048 | 9.6 | 13.1 | qsine:3 | qsine:3 | 107 | 2048 | ^1^H 50W 2.325us ^13^C 80W 4.4us ^15^N 100W 4.5us | 750MHz /  1.9mm HXY | 33.33 kHz / 236K / 270K | 6.4 |

Notes:

The isotopic labeling scheme is denoted as follows: the prefix 'u' indicates uniform labeling with ¹³C, '2' denotes labeling with 2-¹³C-glycerol, and '13' signifies labeling with 1,3-¹³C-glycerol. All labeled samples are uniformly labeled with ¹⁵N.

Acq_td_f1 and Acq_td_f2 represent the number of acquisition points in the time domain for the first and second dimensions, respectively. Acq_time_f1 and Acq_time_f2 are the acquisition times in milliseconds for the first and second dimensions. Proc_f1 and Proc_f2 denote the window functions applied to f1 and f2. For window functions, qsine(t) is defined as: $qsine\left( t \right)=\sin^{2} \left( \frac{t}{AQ}\left( \pi-\phi\right)+\phi\right)$ where $\phi=\frac{\pi}{\mathrm{SSB}}$ , and SSB is set to 3 for qsine-processed spectra. gm(t) function is defined as: $gm\left( t \right)=\exp\left( -at+bt^{2} \right)$ where $a=\pi\cdot LB$ and $b=\frac{a}{2GB\cdot AQ}$ , AQ represents the acquisition time, with GB and LB set to 0.08 and -30, respectively. Proc_td_f1 and Proc_td_f2 are the effective number of points used for processing, where some entries have LP, indicating linear predicted FID appended. The Powers entry indicates the lengths of π/2π/2 pulses for different channels. Experimental conditions include both the set and actual temperatures, with actual temperatures estimated by considering heating from spinning and/or RF. The S/N is the signal-to-noise ratio estimate based on a single resolved peak.

**Table S10.** Table of 3D experiments

| Dataset_ID | Sample | Experiment | Acq_ td_f1 | Acq_ td_f2 | Acq_ td_f3 | Acq_ time_ f1 | Acq_ time_ f2 | Acq_ time_ f3 | Proc_f1 | Proc_f2 | Proc_f3 | Proc_ td_f1 | Proc_ td_f2 | Proc_ td_f3 |
| --- | --- | --- | --- | --- | --- | --- | --- | --- | --- | --- | --- | --- | --- | --- |
| 23 | u_homoM45 | NCOCX | 52 | 132 | 2048 | 3.9 | 4.0 | 15.3 | qsine:3 | qsine:3 | qsine:3 | 52 | 132 | 1024 |
| 24 | uM45_naRIPK3 | CANCO | 70 | 44 | 2048 | 5.3 | 4.6 | 13.1 | qsine:3 | gm:  -30; 0.08 | qsine:3 | 70 | 44 | 1024 |
| 25 | uM45_naRIPK3 | NCACX | 36 | 90 | 2560 | 3.2 | 4.1 | 16.4 | gm:  -30; 0.08 | gm:  -30; 0.08 | gm:  -30; 0.08 | 36 | 90 | 1024 |
| 26 | naM45_uRIPK3 | NCOCX | 30 | 76 | 2048 | 4.1 | 4.1 | 15.4 | qsine:3 | qsine:3 | qsine:3 | 30 | 76 | 1000 |

| Powers | Instrument  (B_s_/ probe) | Experiment  Condition (spinning / set temp / actual temp) | S/N |
| --- | --- | --- | --- |
| ^1^H 220W 3us ^13^C 95W 4.75us ^15^N 150W 5.35us | 900MHz /  3.2mm Efree | 16kHz /  273K / 280K | 9.5 |
| ^1^H 50 W 2.25us  ^13^C 80 W 4.35us ^15^N 100W 4.55us | 750MHz /  1.9mm HXY | 33.33 kHz /  236K / 270K | 3.1 |
| ^1^H 50 W 2.3us  ^13^C 80 W 4.4us ^15^N 100W 4.5us | 750MHz /  1.9mm HXY | 16.66kHz /  265K / 285K | 2.6 |
| ^1^H 240W 2.88us ^13^C 95W 4.25us ^15^N 150W 5.25us | 900MHz /  3.2mm Efree | 16kHz /  273K / 280K | 3.7 |

**Table S11.** NMR and refinement statistics for protein structures

| **NMR distance and dihedral constraints** |  |
| --- | --- |
| Distance constraints | 233 |
| Unambiguous distance constraints | 215 |
| Intra-residue | 0 |
| Inter-residue | 215 |
| Sequential (\|*i* – *j*\| = 1) | 144 |
| Medium-range (\|*i* – *j*\| < 4) | 24 |
| Long-range (\|*i* – *j*\| > 5) | 18 |
| Intermolecular (NMR-based)  Intermolecular (knowledge-based) *  Ambiguous distance constraints | 9  20  16 |
| Hydrogen bonds | 2 |
| Total dihedral angle restraints |  |
| φ | 31 |
| ψ | 31 |
| **Structure statistics** |  |
| Violations (mean and s.d.) |  |
| Distance constraints (Å) | 0 |
| Dihedral angle constraints (º) | 0 |
| Max. dihedral angle violation (º) | 0 |
| Max. distance constraint violation (Å) | 0 |
| Deviations from idealized geometry |  |
| Bond lengths (Å) | 0 |
| Bond angles (º) | 0 |
| Impropers (º) | 0 |
| Average pairwise r.m.s. deviation** (Å) |  |
| Heavy | 1.07 |
| Backbone | 0.52 |

Notes: * :Inter-strand C-C distance constraints on residues with high probability of beta-sheet(	greater than 0.5).

**: 10 structures calculated, average pairwise r.m.s.d based on structure #1.

**Table S12.** Cryo-EM data collection and processing

|  | **Homomeric M45** |
| --- | --- |
| **Sample conditions** |  |
| Buffer | 38.7 mM Acetic acid, 11.3 mM Sodium Acetate, pH 4.0 |
| Assembly method | Dialysis at room temperature overnight |
| Final concentration | 20 µM |
| **Data collection and processing** |  |
| Voltage (KeV) | 300 |
| Electron exposure (e−/Å) | 65 |
| Defocus range | -1.5 to -1.0 |
| Pixel size | 0.824 |
| Micrographs collected | 14,767 |
| Symmetry imposed | C1 |
| Initial particle images (no.) | 457,313 |
| Final particle images (no.) | 11,918 |
| Helical twist (°) | -5.34 |
| Helical rise (Å) | 4.8 |
| FSC threshold | 0.143 |
| Map resolution (Å) | 3.96 |
